# Impact of alternative splicing on Arabidopsis proteome

**DOI:** 10.1101/2024.02.29.582853

**Authors:** Andres V Reyes, Christopher Zhang, Sumudu S Karunadasa, Ruben Shrestha, TaraBryn S Grismer, Danbi Byun, Shou-Ling Xu

## Abstract

Limited proteomic evidence makes it unclear to what extent alternative splicing (AS) isoforms are translated and functionally relevant in eukaryotes. Here, we present a comprehensive proteomic analysis in plants using large-scale data mining, extensive fractionation of AspN-and trypsin-digested proteomes, and both label-free and TMT labeling. In total, we identified 471,196 peptides from 22,479 proteins by searching against Araport11, revealing 32,110 isoform-specific peptides. Using an integrated proteogenomic workflow coupled with SUPPA, we classified these peptides into 2,442 AS events, 879 of which involved intron retention (IR). Further analysis of unannotated events revealed 91 additional IRs that are translated, confirming that retained introns can give rise to peptides. AlphaFold modeling predicted the structural and functional impacts of these isoforms. Our dataset improved existing gene model annotations. By comparing wild-type plants with the AS mutant *acinus pinin*, we found that IR regulates transcript and protein abundance nonlinearly. Phenotypic assays confirmed the functional consequences, including reduced chlorophyll, impaired growth, and increased anthocyanin. Take together, our results provide large-scale proteomic evidence that AS isoforms are translated in plants, demonstrating how AS diversifies the proteome, regulates protein abundance, and affects growth and development.

## Introduction

Alternative splicing (AS) is a widespread and conserved regulatory mechanism, affecting over 95% of multi-exon genes in animals and 70% in plants (1–3). AS is essential for development, tissue identity, immunity, and stress responses across species, and its dysregulation is linked to diseases such as cancer (4–10). Functional studies have highlighted AS’s impact on growth and development across species (11–14). For example, thousands of Dscam1 isoforms are essential for normal neural circuit development in the Drosophila (11,12). Other examples include plant Rubisco activase and *HYPERSENSITIVE TO ABA1* (*HAB1*) which generate isoforms with distinct regulatory roles (15–17), while retained introns in animals can confer growth advantages to cancer cells (18). These examples underscore AS as a key mechanism shaping gene regulation and adaptation across evolution.

Although plants and animals share the same major types of AS events, the frequency and predominant forms differ significantly between the two kingdoms (19,20). Exon skipping is the most common type in metazoans, whereas intron retention is the dominant form in plants, accounting for over 60% of events in plants (3,21,22). The exact reasons for this divergence remain unclear but likely involve a combination of cis-regulatory elements, such as intron length, and differences in splicing factors (20,23).

These differences may have various consequences for the proteome (7,19,22,24,25). Exon skipping and the use of alternative 5′ or 3′ splice sites are thought to increase proteome diversity by creating different protein isoforms or by modulating gene expression through mechanisms such as nonsense-mediated decay (NMD) and uORFs (19,26). Intron retention, is also hypothesized to increase diversity by introducing novel sequences or truncated isoforms. Emerging studies also propose that retained introns may play additional regulatory roles. For example, they may be exported to the cytoplasm for degradation by NMD, sequestered in the nucleus for delayed processing into fully spliced mRNAs (13,27), or degraded by the nuclear exosome. Most of these outcomes, particularly in plants, remain speculative and require experimental validation to determine their actual impact on proteome regulation.

A key unresolved question is the extent to which splice variants are translated and functionally relevant (22,28,29). Evidence from animal studies suggests that AS events often preserve frame and exhibit evolutionary conservation (30,31). Ribosome profiling studies further indicate that many transcript variants associate with polysomes, suggesting that they are translated (32–36). In addition, many splicing events and their regulatory elements are conserved across species, for example, splicing signals in retained introns are found to be more conserved than those constitutive introns (37,38), implying selective pressure to preserve their functions. However, many of these AS variants may be expressed at low levels, leading to debate about their overall functional significance within the proteome.

Direct evidence of splice variant translation at the proteome-wide level has come from mass spectrometry-based analyses. However, the limited number of available studies has yielded conflicting conclusions (39–41), highlighting persistent technical challenges associated with proteomics approaches. These challenges include limited proteomic coverage and sensitivity, the tissue-or condition-specific nature of many AS events, and the difficulty of detecting splice junctions using standard tryptic digestion. This difficulty arises because evolutionarily conserved nucleotide usage at exon boundaries increases the frequency of lysine-and arginine-coding triplets at exon ends, making detection of exon-exon and exon-intron junctions challenging (42). Several efforts have begun to address these barriers in human systems. For example, some studies have leveraged splicing mutants (43), while others have applied deep proteome sequencing strategies (44). However, comparable large-scale efforts remain lacking in plants.

To address this knowledge gap, we conducted a thorough investigation into how AS influences the *Arabidopsis thaliana* proteome. By analyzing over 900 liquid chromatography– tandem mass spectrometry (LC-MS/MS) datasets—720 from public repositories and 180 generated in-house—we achieved the first large-scale identification of isoform-specific peptides in plants. Using our in-house proteogenomic approach, coupled with SUPPA (45) for event classification and additional library annotations, we detected a total of 2,533 splice events. Our results demonstrate that diverse AS events are translated and detectable at the proteome level, substantially expanding the diversity of the *Arabidopsis* proteome. Notably, not only are canonical events such as exon skipping and 5’ or 3’ alternative splicing translated, but many IR events are also translated. This proteomic evidence further provides a better basis for gene model annotation.

We further incorporated the *acinus pinin* double mutant, which has previously been shown to exhibit 2,446 altered AS events, including 1,106 intron retention, 206 exon skipping, 371 alternative donor, 498 alternative acceptor, and 265 combined alternative donor-acceptor events (23,46). Many of these regulated intron retention events can be recapitulated in wild-type plants subjected to biotic or abiotic stress. Our approach allowed us to quantitatively analyze the impact of retained introns on transcripts and proteins, revealing a non-linear relationship between intron retention and transcript/protein abundance. Our data further show that these impacts, driven by both AS and transcriptional regulation, are functionally significant, correlating with the phenotypic defects observed in the *acinus pinin* mutant.

Take together, our findings provide the first large-scale demonstration directly linking AS variation to proteomic outcomes in plants, offering new insights into the regulatory impact of AS on plant development and physiology.

## Results

### Intron retention events detected in *acinus pinin* mutants are recapitulated in wild-type plants under abiotic or biotic stress

Intron retention (IR) is the most prevalent type of AS in plants and is often responsive to stress (19,25). Therefore, we examined whether the altered IR events observed in the *acinus pinin* mutant overlap with stress-responsive events in wild-type plants. To accomplish this, we searched for altered IR events detected in the mutant using *PlantIntronDB* (47) and prioritized those reported in independent studies under similar stress conditions. Several altered IR events present in *acinus pinin* mutants occurred in response to cold, osmotic stress, dehydration, flagellin, fungal infection or heat (Supplementary Fig. S1A-E and Supplemental Table S1).

To evaluate the statistical significance of the observed overlaps, we reanalyzed a few publicly available RNA-seq datasets that met the following stringent criteria: at least three biological replicates; paired-end reads; sufficient sequencing depth; and stress treatments applied at the seedling stage (Supplementary Table S1). We observed significant overlap in IR events between *acinus pinin* mutants and wild-type plants subjected to cold (PRJNA1062790) (48), heat(PRJNA1262915), and flagellin (PRJNA1124505) (49) treatments (Fig. 1A-D, Supplemental Fig.S2A-C). While some IR events were shared across stress conditions, others were stress-specific (Fig. 1A, E-G), highlighting a complex and condition-dependent splicing landscape. Notably, IR events shared between the *acinus pinin* mutant and cold or heat treatments were predominantly upregulated (Fig.1B-C). Conversely, IR events associated with the flagellin response exhibited an opposite pattern of regulation (Fig.1D). However, this trend was not evident at the level of differentially expressed genes (Supplemental Fig. S2D-I).

**Figure 1.**
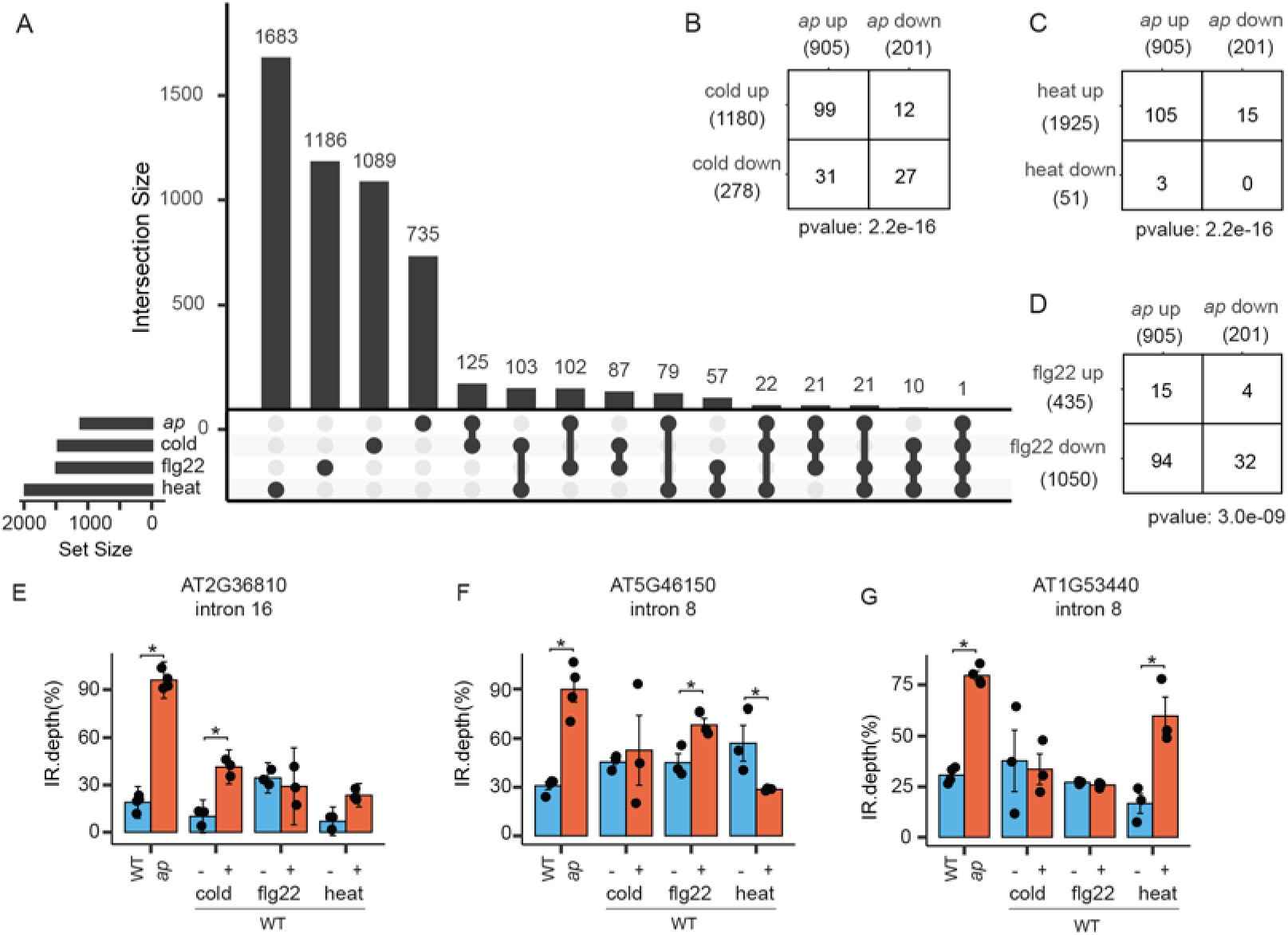
Intron retention events in *acinus pinin* mutants overlap with stress-induced IR events in wild-type plants. (**A**) UpSet diagram showing the overlap of differentially retained introns between the *acinus pinin* mutant and wild-type plants subjected to cold, heat, or flg22 treatment. (**B**–**D**) Pairwise comparisons of up-and down-regulated retained introns in the wild-type upon various stress conditions and in the *acinus pinin* mutant. (**E**–**G**) Representative examples of retained introns. Statistically significant differences (t-test) (p ≤ 0.05) are indicated by an asterisk (*). Error bars represent the mean ± standard deviation.

Together, these results imply that many IR events in the mutant mirror the physiological responses to both abiotic and biotic stress observed in wild-type plants. These findings imply ACINUS and PININ as key integrators of environmental signals in the regulation of AS. We included the *acinus pinin* mutant in our qualitative proteomic analysis to identify the splice variants, and in our quantitative analysis to evaluate the impact of IR on transcript and protein abundance.

### Alternative splicing makes a substantial contribution to the diversity of proteins validated by proteomic analysis

To investigate whether splice variants are translated, we analyzed large-scale proteomics datasets, including data from the *Arabidopsis* draft proteome project (50). This reference dataset, generated from 30 distinct tissues, involved tryptic digestion combined with extensive fractionation and comprised 720 nLC-MS/MS runs. In addition, we generated two complementary datasets: (1) 90 runs from AspN-digested, fractionated proteomes of wild-type and *acinus pinin* mutant seedlings, and (2) 99 trypsin-digested TMT-labeled, fractionated proteome runs derived from pooled peptide samples of wild-type and *acinus pinin* mutants (Supplementary Fig.S3) (see Methods). Together, these datasets encompass 909 nano-scale liquid chromatography-tandem mass spectrometry (nLC-MS/MS) runs (Fig. 2A).

**Figure 2.**
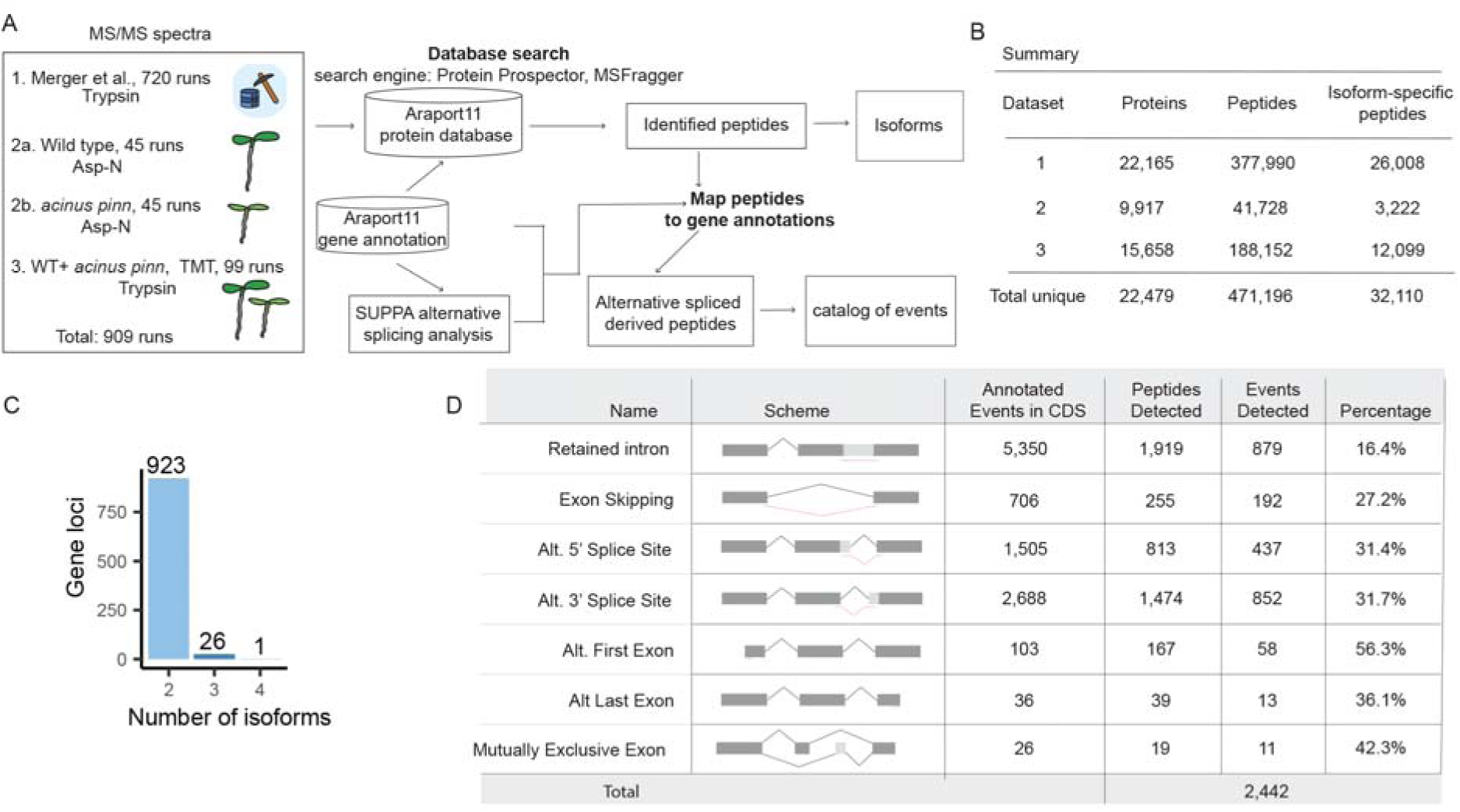
Alternative splicing makes a substantial contribution to the diversity of proteins validated by proteomic analysis. (A) Computational workflow for identifying splice isoform-specific peptides and mapping alternative splicing (AS) events. A total of 909 nanoscale LC-MS/MS runs, including both trypsin-and AspN-digested samples, were analyzed to maximize proteome coverage, enable isoform quantification, and catalog AS events. Proteome-to-genome mapping strategy was performed and integrated with SUPPA-derived splicing coordinates for comprehensive AS annotation. (B) Summary table showing the number of proteins, total peptides, and isoform-specific peptides identified across three datasets, as well as the combined unique totals, highlighting large-scale peptide identification achieved in the *Arabidopsis* proteome. (C) Distribution of gene loci with proteomic evidence supporting 2, 3, or 4 isoforms based on isoform-specific peptides. (D) Summary of AS events annotated in Araport11 and detected by proteomics. Events are grouped by AS type within coding regions (CDS), with peptide identifications supporting each type. Detection percentages represent detected versus annotated events.

All MS Data were acquired using Orbitrap mass spectrometers, which provides high-resolution MS1 acquisition for more accurate identifications. The resulting MS data were searched against the Araport11 protein database (51), which provides a reannotation of the *Arabidopsis thaliana* genome. This more comprehensive database, which is larger than the canonical TAIR annotation, contains 48,354 protein-coding transcript isoforms encoded by 27,650 genes, representing a total of 40,779 non-redundant protein forms. Importantly, 31% of these protein-coding genes in this database encode more than one distinct isoform (Supplementary Fig. S4A), with the majority producing two (Supplementary Fig. S4B). To maximize peptide and protein identification, database searching was performed using two complementary search engines: Protein Prospector and MSFragger (52–54).

Using the *Arabidopsis* draft proteome and our in-house data, we identified a total of 471,196 unique peptides from 22,479 gene loci (Fig.2B). We systematically refined this dataset through a multi-stage filtering pipeline (Supplemental Fig. S5A–B) to isolate peptides specific to particular protein isoforms. The filtering removed 280,916 peptides for single-isoform genes, 12,412 that were not locus-specific, and 145,758 peptides shared among all isoforms. Our final high-confidence dataset thus contained 32,110 peptides that exclusively identify specific isoforms (Fig. 2B, Supplemental Table S2). Subsequent mapping of these isoform-specific peptides (Supplemental Fig. S6A–C) confirmed the expression of multiple protein variants at 950 gene loci (Fig. 2C). This analysis revealed 923 loci with two isoforms, 26 with three, and one with four, confirming a greater number of multi-isoform loci than was previously documented (50).

The classification of AS events and the extent of their translation in plants have remained unknown, despite previous large-scale proteomic studies in *Arabidopsis* (50,55). To investigate this, we developed and employed an integrated proteogenomics workflow with SUPPA event classification (Fig. 2A). Our robust pipeline directly mapped peptides derived from mass spectrometry to their genomic locations in the Araport11 genome, achieving high resolution by accounting for intron-exon junctions and codon ambiguity. Using SUPPA’s splicing coordinates, we accurately classified isoform-specific peptides to identify AS event types (see Methods) (Supplemental Table S2). This analysis identified 2,442 AS events (Fig. 2D), categorized as: 879 intron retentions, 192 exon skippings, 437 alternative 5′ splice sites, 852 alternative 3′ splice sites, 58 alternative first exons, 13 alternative last exons, and 11 mutually exclusive exons. Importantly, this dataset provides the first direct proteomic evidence demonstrating the translation of numerous intron retention events in plants.

### Representative alternative splicing events validated by proteomic evidence

Figure 3 presents representative cases of AS captured at the proteomic level. AspN digestion provided complementary sequence coverage by generating unique peptides that confirmed the presence and translation of distinct spliced isoforms.

**Figure 3.**
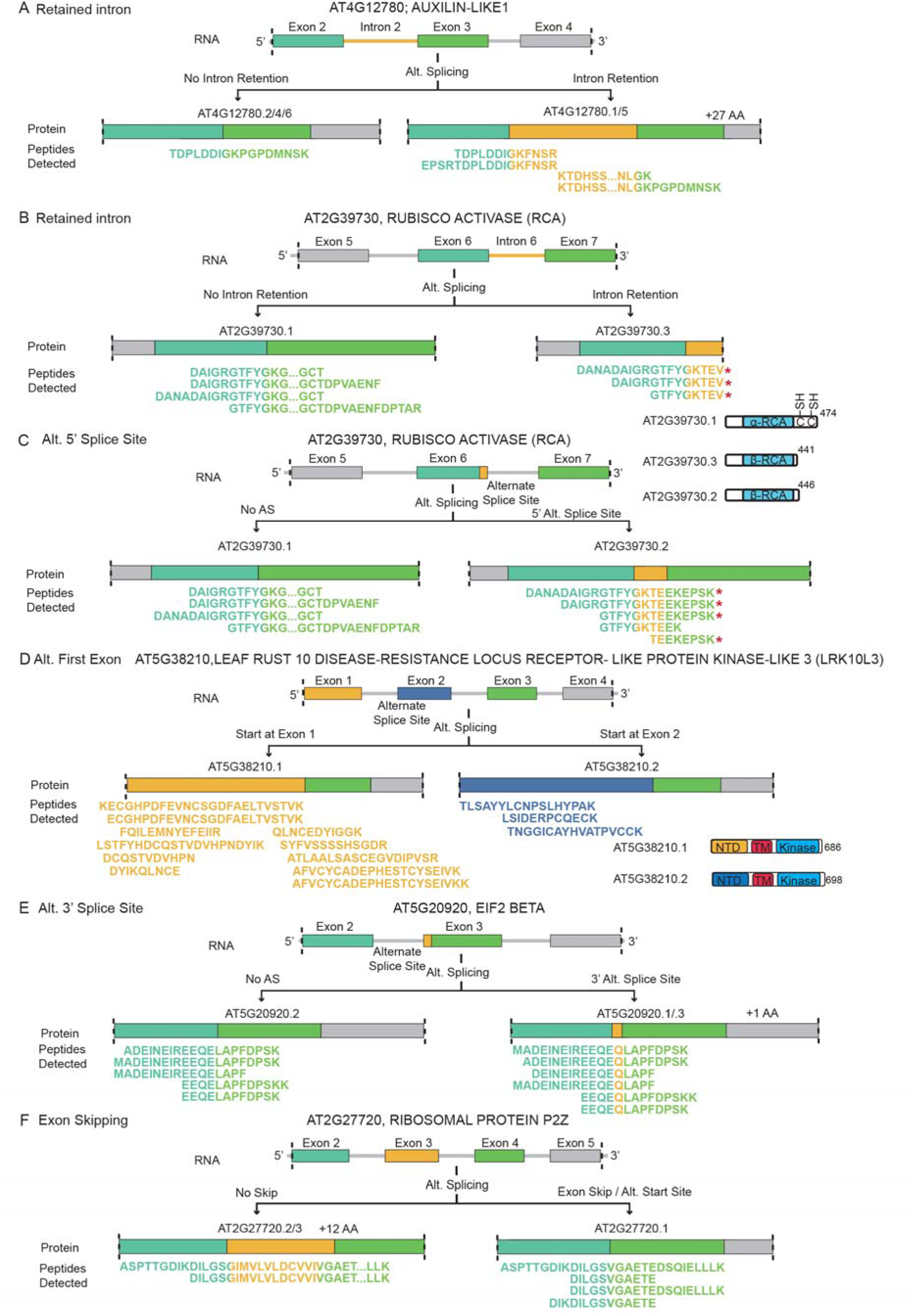
Representative alternative splicing events validated by proteomic evidence. Junction and isoform-specific peptides identified by deep proteomics provide direct evidence for the translation of both isoforms across multiple types of alternative splicing events: intron retention (A–B), alternative 5′ splice site (C), alternative first exon (D), alternative 3′ splice site (E), and exon skipping (F). (**A**) Intron retention in *AUXILIN-LIKE1* generates two isoforms, one of which has a 27 amino acid (aa) extension translated from the retained intron. Both isoforms are supported by peptide evidence. (**B**–**C**) Peptide evidence confirms the translation of three Rubisco activase (RCA) isoforms: a full-length isoform and two truncated isoforms. Intron retention produces a 441aa isoform, while the use of an alternative 5′ splice site introduces a frameshift, yielding a 446 aa isoform. Both truncated isoforms lack redox-sensitive cysteines and are not subject to redox regulation. (**D**) Proteomic support for both isoforms of LRK10L3 demonstrates alternative first exon usage, producing isoforms with distinct N-terminal extracellular domains but identical transmembrane and C-terminal kinase domains. (**E**) Isoform-specific peptides confirm that an alternative 3′ splice site in EIF2 BETA generates two isoforms, one of which carries an additional glutamine residue (Q). (**F**) In ribosomal protein P2Z, peptide evidence validates the translation of both isoforms, with exon skipping producing a shorter isoform lacking 12 amino acids.

In Auxilin-like 1, frame-preserving intron retention (exointron) of intron 2 adds 27 amino acids to the N-terminal intrinsically disordered regions (Fig. 3A, Supplementary Fig. S8A–B). Peptide evidence confirms expression of both the spliced (exon 2–exon 3) and intron-retained (exon 2–intron 2 and intron 2–exon 3) isoforms, with five unique junction peptides identified.

Rubisco activase (RCA; Fig. 3B–C) has been re-examined, revealing a third isoform and revising our understanding of the previously defined RCA-β. While earlier work identified only the full-length RCA-α and a shorter RCA-β (15,16), our proteomic data confirms the translation of three distinct isoforms. Junction peptides provided evidence for the full-length isoform (AT2G39730.1, 474 aa), derived from the exon 6–exon 7 junction, as well as a 441 aa isoform (AT2G39730.3), which results from intron 6 retention (exon 6–intron 6 junction). Furthermore, five peptides confirmed an intermediate 446 aa isoform (AT2G39730.2), generated by a frameshift from an alternative 5’ splice site. Our findings show that the previously considered single RCA-β isoform consists of these two shorter variants, both of which lack the redox-sensitive cysteines and are thus insensitive to redox regulation (Supplementary Fig.S9A) (15,16).

In LEAF RUST 10 DISEASE-RESISTANCE LOCUS RECEPTOR-LIKE PROTEIN KINASE-LIKE 3 (LRK10L3)(56) (Fig. 3D), multiple peptides supported the use of an alternative first exon, producing isoforms with distinct N-terminal extracellular domains but identical transmembrane and C-terminal kinase regions (Supplementary Fig.S9B). This modular arrangement may enable recognition of diverse extracellular signals while retaining a common signaling core.

We also detected an alternative 3′ splice site in EIF2 BETA (Fig. 3E, Supplementary Fig. S10A) that inserts an additional glutamine residue, and exon skipping in ribosomal protein P2Z (Fig. 3F, Supplementary Fig.S10B), further illustrating the diversity of AS events detectable by proteomics.

Collectively, these examples provide direct proteomic evidence that a wide spectrum of AS isoforms are translated *in vivo*. In addition to expanding proteome diversity, many of these isoforms likely alter protein structure and function, as demonstrated by AlphaFold structure predictions, with potential biological consequences.

### Detection of translated retained introns improves annotation of plant proteome

We show that Intron retention, the most prevalent form of AS in plants, contributes to proteome diversity (Fig.2D, Fig.3A-B). While many IR events are captured in the Araport11 annotation (Fig. 2D), a substantial proportion is missing. To systematically assess these unannotated events, we analyzed transcripts with an IR ratio greater than or equal to 15% (i.e., ≥ 15% of transcripts retaining the intron) in wild-type or *acinus pinin* seedlings using paired-end RNA-seq data from seedling tissues (23). This analysis identified 4,931 IR events, of which 2,458 (49.8%) were annotated in Araport11 and 2,473 (50.2%) were absent. Because proteomic searches depend on reference protein databases, the absence of unannotated IR sequences limits the ability to detect AS-derived peptides by mass spectrometry.

To address this limitation, we created an expanded protein database that incorporates unannotated IR events within coding regions (CDS) (Fig. 4A-B, Supplementary Fig. S11). Of the 1,901 unannotated CDS-IR events, 1,742 (91.6%) were predicted to produce truncated proteins, compared with only 509 (40.2%) of the 1,265 annotated IR events in Araport11. Such truncated proteins typically result from premature stop codons or frameshifts introduced by retained introns, producing shorter polypeptides than the fully spliced isoforms.

**Figure 4.**
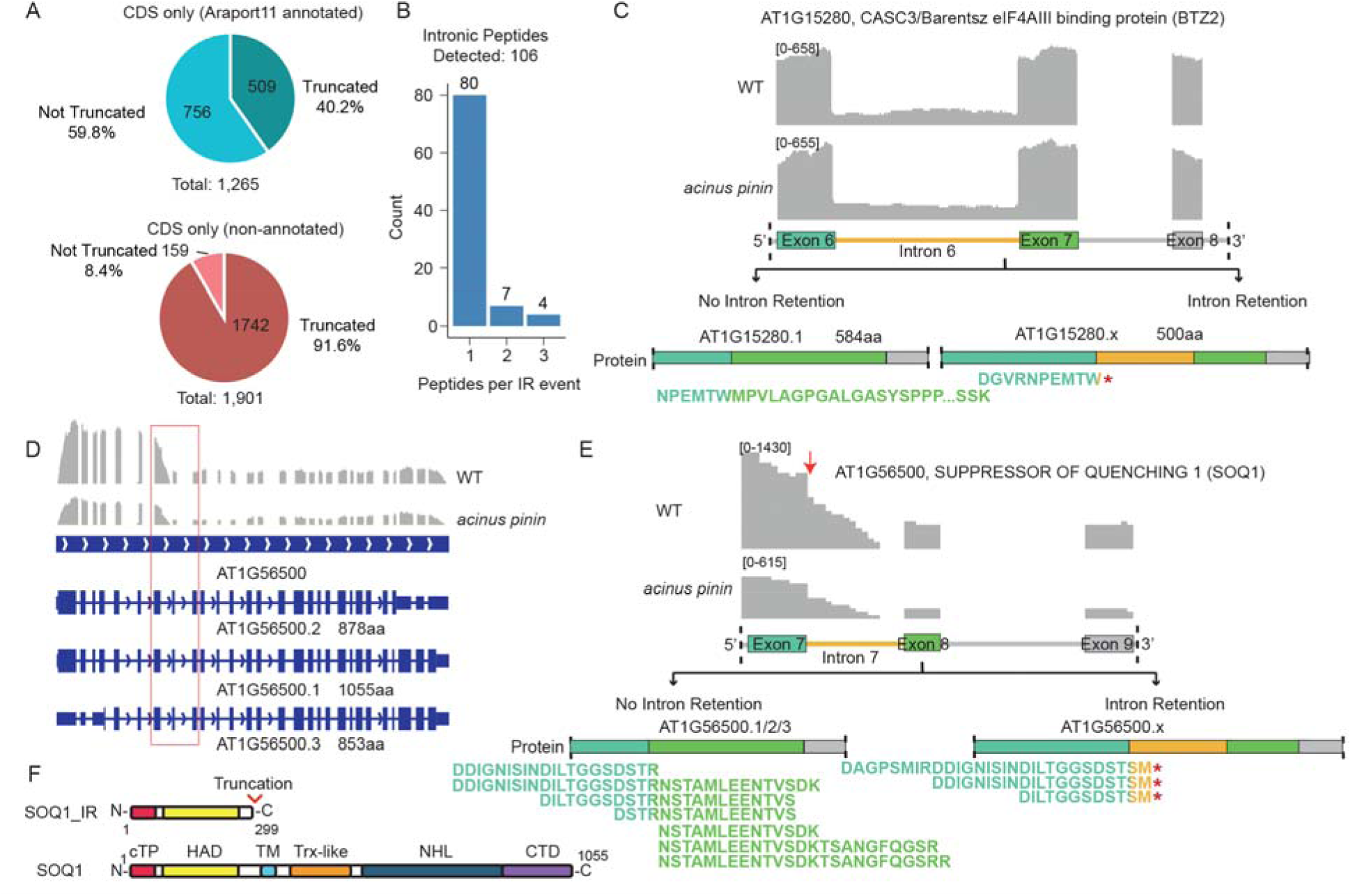
Detection of translated retained introns improves annotation of plant proteome. (A) Comparison of intron retention (IR) effects in annotated versus non-annotated coding sequence (CDS) events. In Araport11 annotations, 40.2% of IR events are predicted to produce truncated proteins, compared with 91.6% of non-annotated events newly identified by RNA-seq. RNA-seq data were obtained from paired-end sequencing of WT and *acinus pinin* mutants (22); events with a retention ratio ≥15% in either genotype were considered. (B) Expanded database in peptide searches identified 106 additional peptides, providing direct proteomic evidence for 91 previously unannotated IR events. (C) Example of truncated protein detection caused by retention of intron 6. Proteomics identified peptides from both the fully spliced and intron-retained isoforms in WT and *acinus pinin* mutants, confirming translation of both. A peptide spanning the exon–intron junction was detected in both WT and mutant backgrounds. (D) RNA-seq and Araport11 annotation of AT1G56500 (SOQ1). While all three annotated isoforms encode transcripts longer than 850 aa, RNA-seq shows a sharp drop in signal across Intron 7 and downstream exons, contradicting the annotation. (E) Proteomic evidence supports both the intron 7–retained truncated isoform and the fully spliced isoform of *SOQ1*. Three junction peptides spanning exon 7–intron 7 confirm the truncated isoform, while four junction peptides spanning exons 7–8 and three peptides from exon 8 support the fully spliced isoform. Retention of intron 7 introduces a stop codon that truncates the protein. (F) The truncated *SOQ1* isoform (SOQ1-IR) lacks the transmembrane, Trx-like, NHL, and CTD domains; the Trx and CTD domains confer redox sensitivity to SOQ1.

We next searched the proteomic data against the combined Araport11 + IR-inclusive database. As noted previously, IR-derived peptides are often challenging to detect because exon-intron junctions are frequently enriched in K/R residues, which can obscure the identification of AS-specific peptides (42,44). Nevertheless, our analysis identified 106 unique peptides supporting 91 previously unannotated IR-derived proteins (Fig. 4B). For example, we detected peptides confirming translation of a truncated isoform (500 aa) of AT1G15280 (CASC3/Barentsz, an eIF4AIII-binding protein, BTZ2), compared to its fully spliced version (584 aa) (Fig. 4C; Supplementary Fig. S12A–B). BTZ2 protein is intrinsically disordered which allows it to interact with various binding partners (57) (Supplementary Fig.S13A). Similarly, peptide evidence supported a truncated isoform (878 aa) of AT4G28650 (PXL2) compared to the full-length protein (1,013 aa) (Supplementary Figure S14).

Integrating proteomics and RNA-seq also allowed us to refine existing genome annotations. For example, although Araport11 predicts three isoforms for AT1G56500 (SOQ1, *SUPPRESSOR OF QUENCHING1*), our data revealed an unannotated IR event in intron 7 that generates a truncated protein of 299 amino acids (Fig. 4D-F; Supplementary Fig. S13B and S15). RNA-seq coverage shows a sharp reduction in read density across intron 7 and downstream exons, while proteomics confirmed translation of the retained intron through three unique peptides, which introduce a stop codon. The data further suggests an alternative 3’end processing within the retained intron. The truncated protein retains the chloroplast transit peptide and HAD domain of the full-length isoform but lacks the transmembrane, Trx-like, NHL, and CTD domains. Notably, the Trx and CTD domains confer redox sensitivity to SOQ1 (Supplementary Fig. S13B) (58,59). In addition, peptide evidence revealed novel isoforms of AT4G31980 (PPPDE thiol peptidase), AT3G50030(ARM-repeat/TPR-like protein), and AT5G02360(DC1 domain-containing protein) (Supplementary Figs S16-S18).

Together, these results demonstrate that combining deep proteomics with RNA-seq uncovers previously hidden IR-derived proteins and refines gene model annotations, thereby expanding the known plant proteome.

### Quantitative TMT proteomics reveals the non-linear impact of intron retention on the proteome

While we detected translation of many retained introns, the overall identification rate remains low, partly because most IR events are only a small proportion of the transcripts. To more comprehensively assess the effects of intron retention, we leveraged the AS mutant *acinus pinin* (*ap*). We performed genome-wide proteomic and transcriptomic profiling of wild-type (WT) and mutants (Fig. 5A), enabling a direct evaluation of how intron retention influences protein abundance.

**Figure 5.**
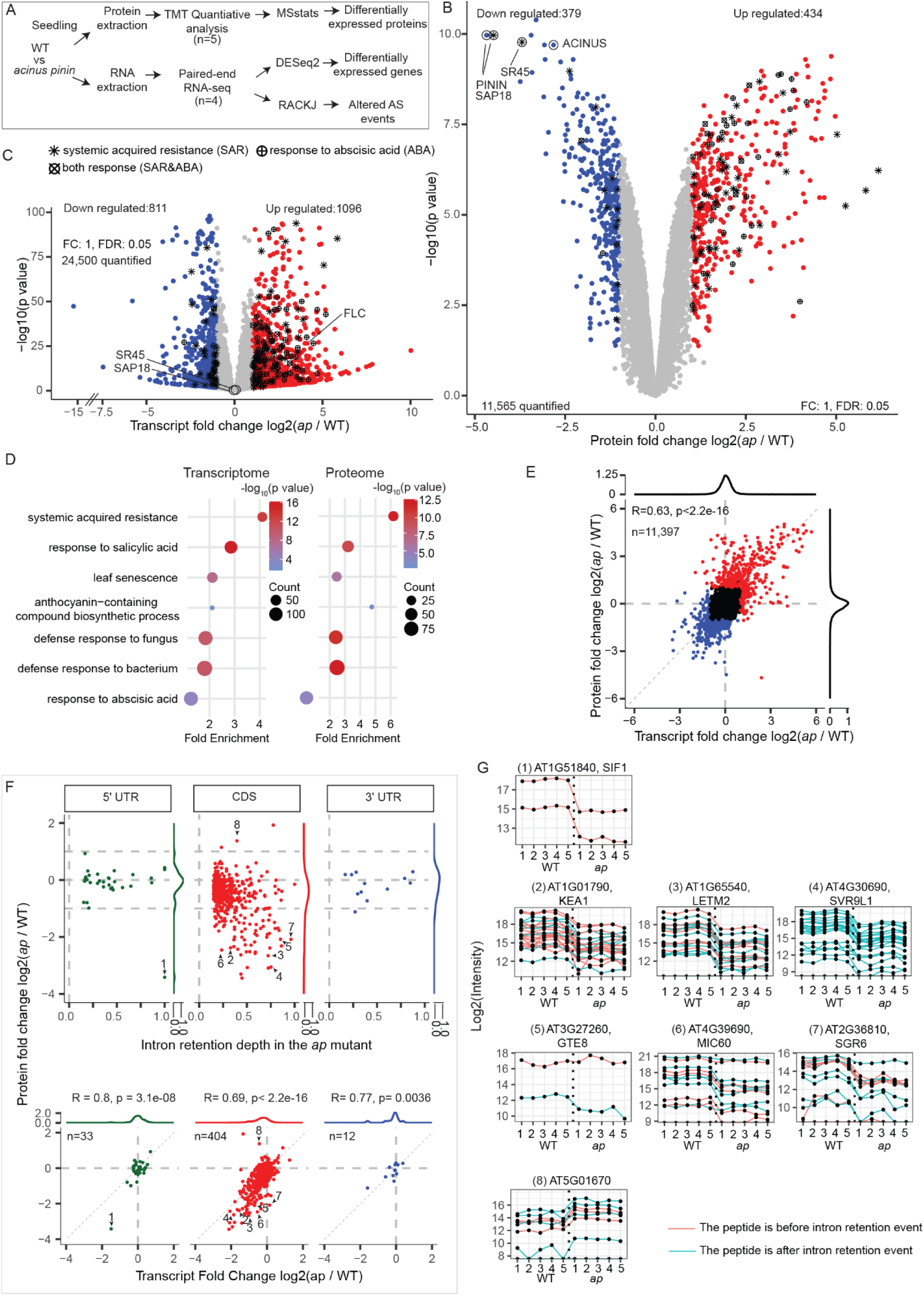
Quantitative TMT proteomics reveals the non-linear impact of intron retention on the proteome (**A**) Genome-wide proteomic and transcriptomic profiling of wild-type and *acinus pinin* mutants. Proteomics was performed with five biological replicates, providing quantitative measurement of protein abundance, while transcriptomics used four biological replicates. (**B**–**C**) Differential expression analysis using TMT-based proteomics and RNA-seq identified hundreds of proteins and transcripts that were up-and down-regulated, including known targets. Proteins and genes associated with systemic acquired resistance and/or ABA signaling are highlighted. Positive controls quantified by TMT (ACINUS, PININ, SR45, and SAP18) and by RNA-seq (FLC) are labeled. (D) Gene Ontology analysis of both proteomic and transcriptomic data revealed additional pathways affected in *acinus pinin* mutants, beyond those reported in our prior study of ABA signaling. (E) Transcript-protein changes show a modest correlation (R=0.63). (F) Nonlinear effects of retained introns that are increased in *acinus pinin* mutants on transcript and protein abundance. Protein abundance changes (y axis) were plotted against RNA changes (x-axis, bottom) and IR depth in the mutant (x-axis, top panel), with density plots also generated. Density plots show the distribution of changes in RNA (bottom panel) and protein (top panel) levels in the *acinus pinin* mutants, both shifted toward negative values for IR events in CDS. Intron retention in untranslated regions (UTRs) had minimal impact on protein levels, except for *AT1G51840* (1). Retained introns in CDS regions typically reduced transcript abundance and caused stronger reductions at the protein level, as illustrated by *AT1G01790/KEA1*, *AT1G65540/LETM2*, and *AT4G30690/SVR9L1* (2–4). Some genes, e.g., *AT3G27260/GTE8*, *AT4G39690/MIC60*, and *AT2G36810/SGR6* (5–7), showed minimal transcript changes but substantial protein decrases. Conversely, IR occasionally increased protein levels without affecting transcripts, as observed for *AT5G01670* (8). (G) Peptide quantifications for the proteins highlighted in (F). Peptides upstream and downstream of retained introns are colored green and red, respectively.

TMT quantified 11,565 proteins (13,850 protein groups) across five biological replicates (Supplementary Fig. S19), while RNA-seq measured 24,500 genes. Differential expression analysis identified hundreds of up-and down-regulated proteins and transcripts, including known targets such as ACINUS, PININ, SR45, and SAP18 at the protein level, and FLC at the transcript level (Fig. 5B–C). Integrative pathway analysis further showed that ACINUS and PININ regulate broad biological processes—including systemic acquired resistance, salicylic acid signaling, senescence, and anthocyanin biosynthesis—beyond their established role in ABA signaling (Fig. 5D). Transcript and protein changes were modestly correlated (Pearson R = 0.63, p < 2.2 × 10⁻¹⁶), indicating widespread post-transcriptional regulation (Fig. 5E).

We then quantified increased IR (IIR) events in *acinus pinin* mutants, defined as ≥15% intron retention depth and ≥1.5-fold change, and assessed their impact on RNA and protein abundance (Fig. 5F). These IIRs occurred in 5′ UTRs (33 genes), CDS regions (404 genes), and 3′ UTRs (12 genes), in contrast to animal systems where IR is enriched in UTRs and non-coding RNAs (24,38). To visualize their impact, protein abundance changes (y axis) were plotted against RNA changes (x-axis, bottom) and IR depth in the mutant (x-axis, top panel), with density plots also generated. Intron retention in UTRs generally did not alter protein levels, except for SIF1 (At1G51840), where IR caused by a marked reduction (Fig. 5F–G). In contrast, CDS intron retention typically led to decreased transcript abundance and an even stronger reduction in protein abundance (e.g., KEA1, LETM2, SVR9L1; numbered 2–4), consistent with NMD-mediated degradation (Fig. 5F-G).

The analysis also revealed non-linear behaviors (Fig.5F-G): some genes showed little change in RNA but strong reductions in protein abundance (e.g., GTE8, MIC60, SGR6), numbered (5–7), while others exhibited protein increases (e.g., AT5G01670; numbered 8). These patterns suggest that IR can alter proteome output through mechanisms beyond simple transcript degradation.

To further investigate, we annotated peptides relative to retained introns, reasoning that truncated proteins would generate differential peptide abundance upstream versus downstream of the IR site. For most genes, peptide levels changed concordantly in the mutant, suggesting that truncated proteins are rarely produced, unstable, or that current TMT methods lack the resolution or precision to detect such differences. In one isolated case (AT3G27260/GTE8), the detected upstream peptide remained unchanged while the downstream peptide was reduced, consistent with the production of a truncated protein, although such instances appear to be rare.

Overall, the effects of IR were non-linear: CDS intron retention strongly reduced mRNA and protein levels, UTR intron retention was largely neutral, and a subset of cases showed disproportionate effects at the protein level. These results indicate that IR exerts complex, context-dependent effects on gene expression and proteome output in Arabidopsis. For events with decreased intron retention, conclusions remain limited, likely due to the smaller number of detectable cases (Supplementary Fig. S20).

### Cumulative effects of intron retention on protein abundance drive mutant phenotypes

The functional impact of alternative splicing on individual proteins is often difficult to assess, partly due to redundancy within protein families and the challenge of knocking out a single isoform without affecting others. However, the striking phenotype of the *acinus pinin* mutant suggests that the altered alternative splicing and transcript change have functional consequences. We therefore tested whether the cumulative downregulation of multiple proteins contributes to *acinus pinin* mutant phenotypes. Analysis of proteomic and transcriptomic data (Fig.6A-B) revealed that STRESS-RESPONSIVE PROTEIN 9-LIKE 1 (SVR9L1) (60), LEUCINE ZIPPER-EF-HAND-CONTAINING TRANSMEMBRANE PROTEIN 2 (LETM2) (61), NIEMANN-PICK DISEASE TYPE C1-2 (ATNPC1-2 (62)), POTASSIUM EFFLUX ANTIPORTER 1 (KEA1), LIMIT DEXTRINASE/ATLDA(63), and SODIUM/HYDROGEN ANTIPORTER 5 (NHX5) (64) were downregulated at both transcript and protein level due to significant intron retention, whereas BRANCHING ENZYME 3 (BE3) (65) and S-ADENOSYLMETHIONINE TRANSPORTER 1 (SAMT1) (66) were downregulated at transcript and protein levels without affecting the splicing. Notably, these genes have been previously reported to influence chlorophyll content and root growth when combined with additional mutations.

**Figure 6.**
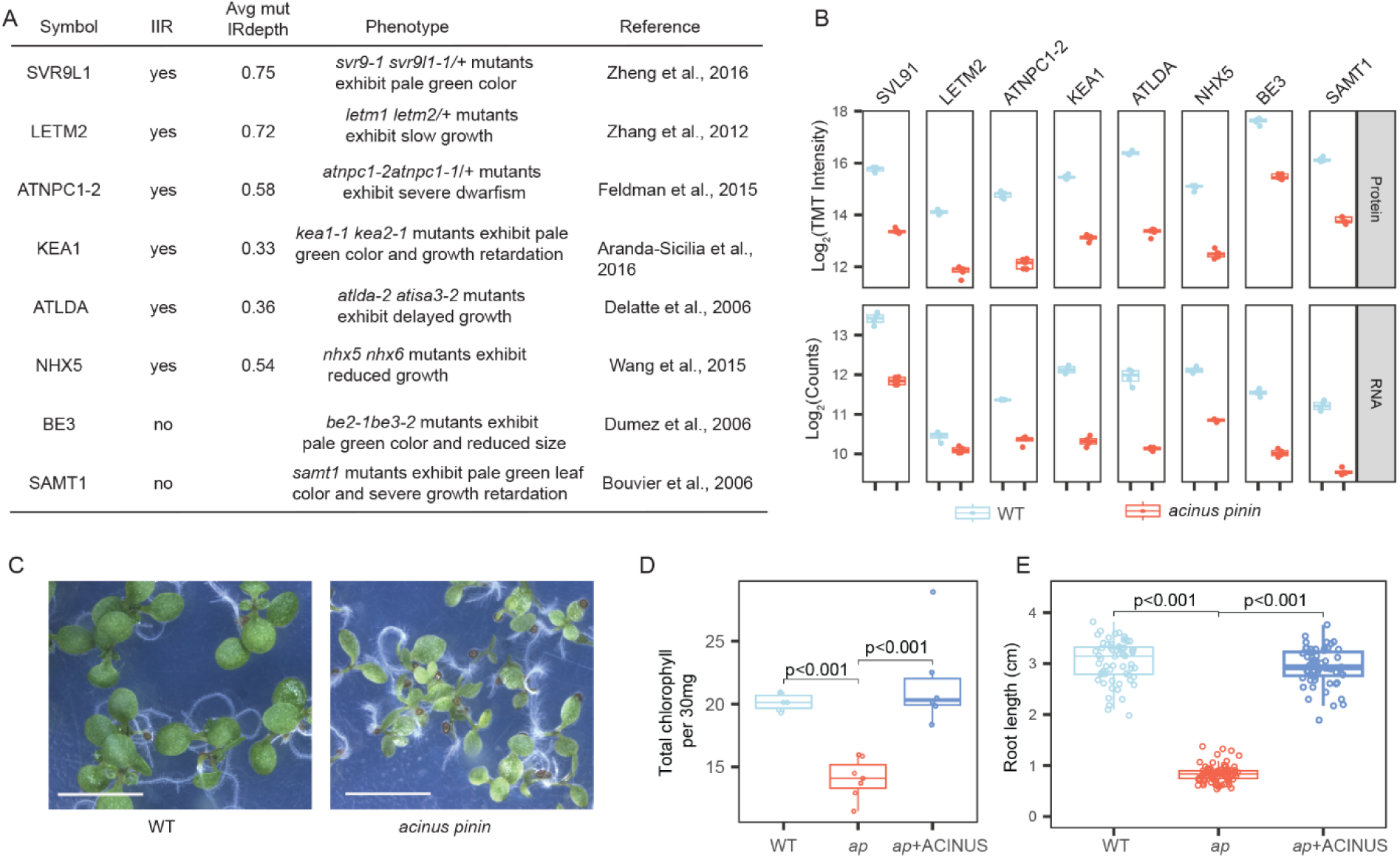
Cumulative effects of intron retention on protein abundance drive mutant phenotypes. (A) Proteins involved in regulating leaf color and plant growth. For genes with increased intron retention (IIR), the average IR depth in the *acinus pinin* mutant is included. (B) Corresponding reductions in protein and transcript levels for these genes, as revealed by quantitative proteomics (log_2_ TMT intensity) and RNA-seq of (log_2_ counts). (C) Phenotype of *acinus pinin* mutants showing reduced growth and pale-green leaves. Scale bar = 2 mm. (**D**–**E**) Quantitative analysis of chlorophyll content (D) and root length (E) in WT, *acinus pinin* mutants, and the complemented line (+ACINUS). Statistical significance was assessed using Student’s t-test.

Guided by these proteomic insights, we measured chlorophyll content and root length, finding significant reductions in both traits in *acinus pinin* mutants that were rescued by ACINUS expression (Fig. 6C–E). These results indicate that the pleiotropic defects may arise from the combined reduction of multiple proteins through IR and other regulatory mechanisms, linking alternative splicing directly to functional outcomes.

### Upregulated proteins in anthocyanin biosynthesis correlate with increased anthocyanin accumulation in *acinus pinin* mutants

Many splicing factors have been reported to influence transcription, although the underlying mechanisms remain unclear (43,67). Guided by transcriptomic and proteomic data, we found that ACINUS and PININ regulate the expression of numerous genes (Supplementary Fig. S2, Supplementary Table S1) without affecting their alternative splicing, indicating regulation at the transcriptional level. Notably, several genes encoding proteins involved in anthocyanin-containing compound biosynthesis, as cataloged by Gene Ontology analysis, were strongly upregulated in the mutants at both the transcript and protein levels (Fig. 5D, Fig. 7A, Supplementary Table S1).

**Figure 7.**
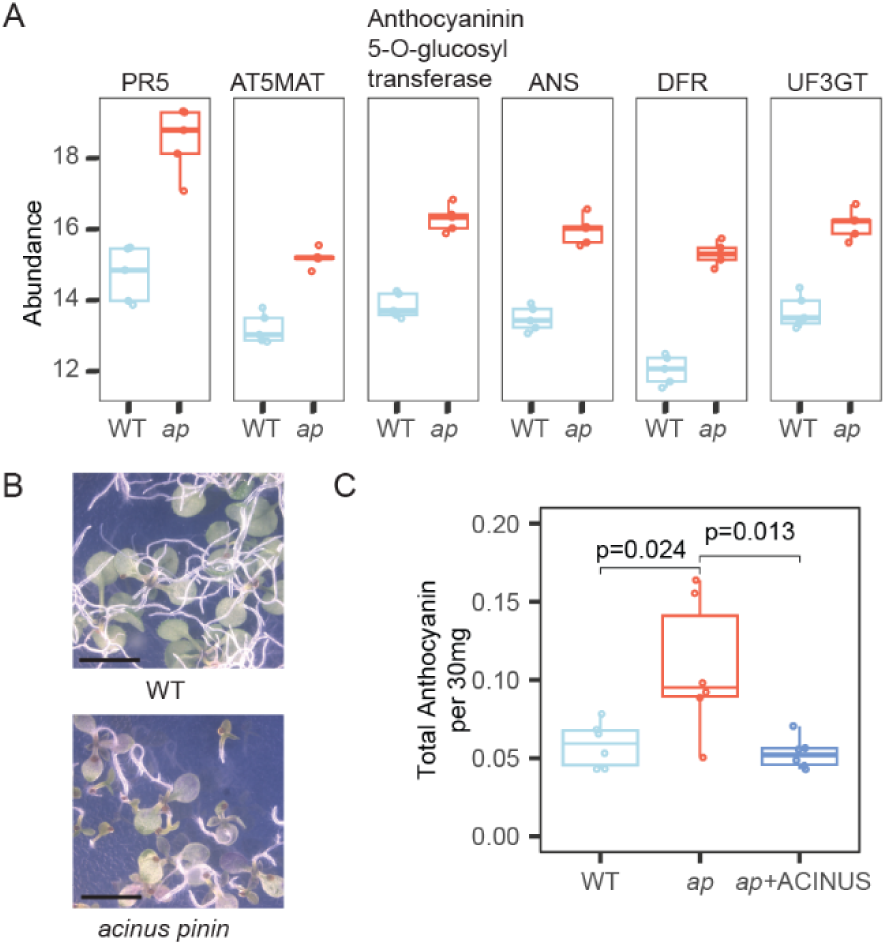
Upregulated proteins in anthocyanin biosynthesis correlate with increased anthocyanin accumulation in *acinus pinin* mutants. (A) Box plot showing the abundance of proteins associated with anthocyanin biosynthesis, as defined by GO analysis, in *acinus pinin* mutants (*ap*, red) and wild type (WT, blue). Protein levels are significantly higher in the mutant. (B) Abaxial cotyledon images showing elevated anthocyanin accumulation in *acinus pinin* mutants. Scale bar, 2 mm. (C) Quantification of total anthocyanin content in WT, *acinus pinin* mutants, and the complemented line (+ACINUS), showing increased levels in the double mutant. Statistical significance was assessed using student’s t-test.

Based on these molecular changes, we predicted altered anthocyanin accumulation in the double mutants and examined their anthocyanin phenotype. Indeed, when grown on ½ MS plates, the mutants exhibited pronounced anthocyanin accumulation beneath the cotyledons, resulting in a distinct purple coloration and quantitatively higher anthocyanin content, both of which were fully rescued by ACINUS expression (Fig. 7B–C). Because both transcript and protein levels are altered, we conclude that the anthocyanin phenotype in *acinus pinin* mutants is driven by transcriptional regulation. Whether ACINUS and PININ directly bind these loci to control expression remains unknown and will be addressed in future studies.

## Discussion

For decades, researchers have debated whether AS represents an evolutionary innovation or merely splicing noise. A central question is whether splice variants are translated and functionally relevant (28,29). Ribosome profiling studies confirm that many transcript variants are associated with polysomes (32–34,36,68), yet it remains unclear whether these translation events yield stable, functional proteins. Proteomics offers more direct evidence, but technical limitations in coverage and sensitivity have constrained conclusions (42), and previous studies have reported conflicting results (39–41). In this study, we analyzed over 900 LC-MS/MS datasets—720 from public repositories and 189 generated in-house—using a custom-expanded library, dual protease digestion (trypsin and AspN), both label-free and TMT labeling, and two complementary search engines (Protein Prospector and MSFragger) (Fig. 2 and 3). This strategy enabled the first large-scale identification of isoform-specific peptides in plants and the identification of 2,533 splice events, demonstrating that diverse AS isoforms, particularly intron retention, are translated and detectable at the proteome level. Although our dataset is smaller in scope than comparable human studies (44), it provides strong evidence that AS substantially expands the Arabidopsis proteome.

Our proteomics data also contributes to improving genome annotation. With or without RNA-seq assistance, we demonstrated that proteomics-based evidence enhances current protein annotations (Fig. 4; Supplemental Fig. S14-S18). Because shotgun proteomics relies heavily on the quality of reference databases, these improvements will directly facilitate future proteomic studies. Further integration of RNA-seq data—particularly full-length sequencing approaches such as Nanopore or PAC-seq—will greatly assist in refining gene models. Continued expansion of the protein search library, informed by these rounds of integration, will further refine the database and enhance the accuracy of proteome-wide analyses.

In addition to expanding proteome complexity by generating multiple distinct protein isoforms, our results demonstrate that alternative splicing (AS) can effectively downregulate gene expression (Fig. 5F), as evidenced by intron retention (IR) in *acinus pinin* mutants. Previous studies in plants have suggested that IR can modulate gene expression via nonsense-mediated decay (NMD), either through analyses of Arabidopsis NMD factor mutants (*upf1-5*, *upf3-1*) or bioinformatic assessments of IR-containing premature stop codons (25,69–71). Here, we provide the first integrated experimental evidence combining RNA-seq and proteomics to show that retained introns within coding sequences not only reduce transcript abundance but often result in even more pronounced reductions at the protein level. Strikingly, for a subset of genes, transcript levels were largely unchanged while protein abundance decreased substantially, highlighting the non-linear and post-transcriptional impact of IR on the proteome. These findings underscore the dual role of AS in both diversifying protein isoforms and fine-tuning gene expression through mechanisms beyond transcript regulation alone.

We further demonstrate that the cumulative effects of multiple AS events contribute to the observed reduced growth and altered chloroplast phenotypes. Many intron retention events are induced in wild-type plants during developmental stage transitions (13) or in response to environmental cues, suggesting that they likely have functional consequences. While establishing the functional relevance of individual AS events has been challenging, limited cases have shown that distinct isoforms can produce proteins with dramatically different properties (72), including dominant-negative truncated forms or proteins that escape the same regulatory mechanisms, as exemplified by RCA proteins. Importantly, our large-scale identification of AS events at the proteome level provides a comprehensive resource that will guide future studies aimed at understanding the functional impact of alternative splicing in plants.

Although ACINUS functions as a splicing factor and integrates stress-responsive signals (Fig. 1), our data show it also regulates gene expression at the transcriptional level (Figs. 5 and 7). In the *acinus pinin* mutant, some phenotypes are driven by transcriptional changes, revealing a non-splicing role for ACINUS. While it directly regulates FLC (46), whether ACINUS controls additional genes directly or indirectly, and whether it acts within the splicing complex or through separate transcriptional machinery, remains an open question for future studies.

## Materials and methods

### Plant materials and growth conditions

Arabidopsis thaliana Col-0 (WT), *acinus-2 pinin-1* (*ap*) mutants, and complementation line (*ap*+ACINUS) (23,46) were used. Seeds were surface-sterilized, stratified for 3 days, and germinated on ½ MS agar plates at 22 °C under constant light. Seedlings were harvested at specific stages. All plates were grown vertically except for Fig. 6C.

### Chlorophyll measurement and root length assay

Chlorophyll was extracted from 30 mg of 7-day-old seedling tissue, following protocol described in (73). Samples were flash-frozen, ground, and extracted with 80% acetone overnight at 4 °C.

After centrifugation (5,544 x g, 15 min, RT), absorbance was measured at 663 nm and 645 nm (Tecan plate reader). Chlorophyll content was calculated as: Chlorophyll = 8.02 × (chlorophyll a absorbance A663) + 20.20 × (chlorophyll b absorbance A645). For root assays, seedlings grown vertically for 7 days on ½ MS were photographed. Images were analyzed in ImageJ, and root lengths were measured from the hypocotyl-root junction to the root tip (WT: n=60; *ap*: n=82; *ap*+ACINUS: n=60). Statistical significance was assessed using Student’s t-test.

### Anthocyanin analysis

Anthocyanin was extracted from 30 mg 11-day-old vertical plates, frozen, ground tissue in 1 mL of 1% HCl-methanol. Samples were vortexed and incubated overnight at 4 °C in the dark, then centrifuged (16,000 x g, 5 min). Absorbance of the supernatant was measured at 530 nm and 657 nm (Tecan plate reader), and anthocyanin content were calculated as (absorbance at 530 (A530)-0.25 x absorbance at 657 (A657)).

### Sample preparation for AspN and TMT11 experiments

Tissue (100 mg) from 9-day-old wild-type and *acinus pinin* seedlings grown on vertical plates was harvested, flash-frozen, and ground. Proteins were extracted as described in (74), with a slight modification to extraction buffer Y (100 mM Tris-HCl, pH 8.0; 2% SDS w/v; 5 mM EGTA; 10 mM EDTA; 1 mM PMSF; 2× protease inhibitor, Roche). Extracts were reduced with TCEP, alkylated, and digested with trypsin, followed by desalting using Sep-Pak columns (Waters). For TMT experiments, five biological replicates per group were prepared.

### TMT 11-plex labeling

TMT 11-plex reagents (ThermoFisher, Cat# A34808) were reconstituted in anhydrous acetonitrile (0.01 mg/μL). Approximately 200 μg of peptides from each sample were dissolved in 66 μL of 50 mM HEPES buffer. Of this, 60 μL (182 μg) was labeled with 40 μL of TMT reagents (TMT126– 130N), while the remaining 6 μL (18.2 μg) was pooled to 60 μL and labeled with 40 μL TMT131C. Reactions proceeded for 1 h at 25 °C with shaking at 700 rpm and were quenched with 5 μL of 1 M Tris (pH 8). Peptides were then acidified with 45 μL of 10% FA in 10% acetonitrile. The final reaction contained TMT at 4 μg/μL, peptides at 2 μg/μL, and 40% acetonitrile (v/v), similar to formular described in (75). The labeling scheme is shown in Supplementary Fig.S3. One-eighth of the 11 labeled samples was combined for high-pH reversed-phase HPLC fractionation (74).

### Fractionation of peptides samples for AspN and TMT samples

High pH reverse-phase chromatography was performed on the Vanquish Flex system (ThermoFisher) equipped with a 4.6×150-mm Gemini 5μm C18 column for TMT-labeled peptides and AspN digested samples. Peptides were loaded onto the column in 25 μL of buffer A (20 mM ammonium formate, pH 10). Buffer B consisted of buffer A with 90% (vol/vol) acetonitrile. Peptides were separated at 0.5 mL/min using high-pH reversed-phase gradients. Common steps included column equilibration at 1% B and sequential increases to 50–70% B, 70–100% B, and 100% B hold. Fraction were collected every 1–2 min. TMT batch 1 and AspN: 5 min equilibration; 1–5% B in 2 min, 5–50% B in 58 min; fractions every 1 min. TMT batch 2: 10 min equilibration; 1–10% B in 10 min, 10–50% B in 40 min; fractions every 2 min. TMT Batch 3: 10 min equilibration; 1–10% B in 20 min, 10–50% B in 80 min; fractions every 1 min. Collected fractions were vacuum-dried and subjected to LC/MS/MS analysis.

### LC/MS/MS analysis of AspN and TMT samples

Peptides were analyzed on a Q-Exactive HF or Orbitrap Eclipse hybrid quadrupole-Orbitrap mass spectrometer (Thermo Fisher) coupled to an Easy LC 1200 UPLC system. Peptides were trapped on an Acclaim PepMap 100 C18 column (75 μm × 2 cm) and separated on a 25 cm × 75 μm, 1.7 μm C18 Aurora column (IonOpticks) at 300 nL/min using a 120 min gradient: 3–28% solvent B (80% ACN, 0.1% FA) over 106 min, 28–44% B over 15 min, followed by a 9 min wash at 90% B. Batch 1 TMT samples were analyzed on the Q-Exactive HF with MS1 scans over m/z 375– 1600 and were acquired at m/z 375–1600, resolution 120,000, AGC 3E6, max injection 100 ms. MS2 scans select the top 20 multiply charged precursors for HCD fragmentation (NCE 30, isolation 1.0 m/z) with a scan range of 200-2000 m/z, resolution 30,000, AGC 5E4, max injection 60 ms, and dynamic exclusion 24 s. Batch 2 and 3 TMT and AspN samples were analyzed on the Eclipse; AspN MS1 scans were acquired at m/z 375–1600, resolution 120,000, AGC 2E5, max injection 50 ms, and MS2 targeted top multiply charged precursors (charge 2–8) with HCD 27%, isolation 1.4 m/z, resolution 15,000, AGC 5E4, max injection 22 ms, cycle time 3 s, dynamic exclusion 30 s. TMT samples were measured using real-time library search (76), with MS1 m/z 400–1600, resolution 120,000, intensity threshold 5E3, charge 2–6, dynamic exclusion 45 s; MS2 in the ion trap (isolation 0.7 m/z, CID 35%, max injection 35 ms); MS3 using synchronous precursor selection of 10 precursors (isolation 0.7 m/z, HCD 65%, Orbitrap resolution 50,000, scan range 100–500 m/z, max injection 200 ms, cycle time 2.5 s). Real-time data searches (77) were performed against the TAIR10 database with carbamidomethyl and TMT11plex (static) and oxidation (variable) modifications.

### Overlap of intron retention analysis by PlanIntrontDB

Introns exhibiting increased retention in *acinus pinin* mutants (single-end RNA-seq) (46) were retrieved from the Plant Intron Splicing Efficiency Database (PlantIntronDB; plantintron.com) (3), prioritizing datasets generated under comparable conditions by independent groups, and used for statistical analysis in Supplemental Fig. S1.

### Data analysis of overlapping differentially expressed genes and altered retained introns

RNA-seq data for wild-type and *acinus pinin* mutants were processed as described (23). Additional RNA-seq datasets for selected biotic and abiotic stresses—flg22 (PRJNA1124505) (49), cold (12 h, PRJNA1062790) (48), and heat (PRJNA1262915)—were downloaded using SRA-Tools (v2.10.0). Reads were aligned with STAR (v2.7.10b) using quantMode=GeneCounts, outFilterMultimapNmax=20, and outSAMattributes=NH HI AS nM MD. Transcriptomic and alternative splicing analyses followed (23,46). Dataset overlaps were evaluated with a hypergeometric test. Pairwise overlaps were classified as concordant (shared up-or downregulated events) or discordant (oppositely regulated), and statistical significance was assessed via Fisher’s exact test on a collapsed 2×2 contingency table. Intron numbers from RackJ analysis (Supplementary Table S1) are assigned relative to the plus strand, from 5′ to 3′; for genes on the negative strand, introns are numbered in descending order. In figures, intron numbers have been adjusted to ascending order for clarity.

### Data search and detection of isoform-specific peptides

Isoform-specific peptides were identified by searching all 909 runs (Fig. 2) using Protein Prospector and MSFragger with similar parameters. Previous studies showed that these two search engines yield complementary peptide identification results (54). For all HCD data, the precursor mass tolerance was set to 10 ppm and MS/MS tolerance to 20 ppm. For CID TMT data, low resolution, precursor mass tolerance was set to 10 ppm and MS/MS tolerance to 0.6 Da. Carbamidomethylation of cysteine was set as a fixed modification, while variable modifications included protein N-terminal acetylation, peptide N-terminal Gln conversion to pyroglutamate, and methionine oxidation. Additionally, for TMT data, a mass of the TMT11 tag (229.16293) was set as a variable modification on lysines or on the N-terminal of a peptide. False discovery rates (FDR) were set at 5% for proteins and 1% for peptides. Cleavage specificity was defined according to the protease used (Trypsin or AspN), allowing one missed cleavage and up to two modifications per peptide. Raw data were searched against the Araport11 protein database, which contains 48,354 entries.

### Peptide-level processing and isoform assignment

Peptide-level identification results were imported into R and processed as follows. Decoy matches were removed, and sequences with N-terminal methionine loss were adjusted accordingly. Peptides from both Protein Prospector and MSFragger searches were combined, retaining only one peptide per distinct sequence. Peptides mapping to single-isoform genes or to multiple genes were excluded. Isoform-specific peptides were defined as those mapping to isoforms disjoint from at least one set of isoforms represented by other peptides from the same gene locus (see algorithm breakdown, Supplemental Fig. S7).

### Proteome-genomic mapping and classification of splicing events

Peptides identified from the database were filtered to retain only those corresponding to genes with multiple distinct protein isoforms. For each peptide, an in-house R script was used to retrieve its position on the protein and the transcript to which it maps. Genomic coordinates of each transcript exon were then obtained. Coding sequence (CDS) exon coordinates were enumerated, and the base pairs corresponding to each amino acid of the peptide were calculated (first amino acid: position ×3 −2; last amino acid: position ×3). Peptides spanning exon junctions were identified by comparing these coordinates with exon boundaries.

In parallel, Araport11 annotations were provided to SUPPA (v2.3) (45) to generate a comprehensive set of annotated alternative splicing (AS) events with genomic coordinates. SUPPA was run under default settings, with the variable option enabled for intron retention events. Peptide-genome coordinates were integrated with SUPPA-derived AS coordinates to identify splicing events. For retained introns, alternative 5′ and 3′ splice sites, and alternative first and last exons, events were called based on coordinate overlap. For exon skipping and mutually exclusive exon events, junction-spanning peptides were used to confirm splicing patterns. All analyses, except for those using SUPPA, were performed with custom R scripts, available in the project repository.

### Creation of custom databases and data search

A custom database of unannotated intron retention (IR) events was generated by first filtering introns with high IR depth ratios (≥0.15) in either wild-type or mutant samples from our high-coverage RNA-seq dataset. Genomic coordinates of the selected introns were retrieved, and introns already annotated in Araport11 (as determined by SUPPA) were excluded to avoid redundancy. Each intron of interest was mapped onto the genomic coordinates of all transcripts for its corresponding gene. Introns fitting without gaps—either immediately upstream of the first exon, downstream of the last exon, or between two exons—were used to assemble the genomic sequences, sorted by strand, using the IRanges R package. The underlying genomic sequences were translated into protein sequences using the BSgenome R package. Newly constructed proteins were appended to the Araport11 protein database for downstream peptide searches. Subsequent searches were performed as described above, retaining only peptides that mapped to the custom proteins.

### Quantitative TMT11 plex data analysis

For Eclipse MS data, raw files were searched using MaxQuant v2.4.2.0. In the global parameters, the type was set to MS3 ion reporter mode using the correction factors provided by the TMT reagents. No normalization setting was set. The first peptide mass tolerance was set to 20 ppm and the main search peptide mass tolerance was set to 4.6 ppm. The MS2 match tolerance was set to 20 ppm. Reporter mass tolerance was set to 0.003, and the isobaric weight exponent was set to 0.75. The reference protein database used was a modified version of the Arabidopsis Thaliana TAIR 10 database, downloaded from TAIR: “TAIR10_pep_20101214.fa”. All other parameters remained at their default. *For protein level analysis*: MaxQuant outputs proteinGroups.txt and evidence.txt were used with R package MStatsTMT (78). The parameters for MSstatsTMT’s proteinSummarization function: method was set to msstats, global normalization was set to true, reference normalization was set to true, and remove normalization channel was set to true. Protein level quantifications for each sample were analysed to impute protein quantifications that were only completely missing in one condition while being completely quantified in all 5 replicates in the other condition. The imputation was done from a normal distribution with a width of 0.3 and downshifted 1.8 from the values of the total matrix. Following imputation, a moderated statistics test was done using MSstatsTMT’s groupComparisonTMT function. Proteins with missing p-values were filtered out. If an ambiguous protein group existed with different gene loci, they were discarded. If multiple protein groups existed for a single gene locus, the protein group with the most peptides quantified was kept (also see Supplementary Fig. 16). Protein groups with a p-value less than 0.05 and a fold change greater than 1 or less than - 1 were considered as differentially abundant proteins. Peptide level quantifications shown in Fig. 5G were generated from Maxquants’s peptides.txt output. To determine each peptide’s position relative to intronic regions, we developed a custom R script that maps intron coordinates onto specific isoforms to retrieve the amino acid position of the intron insertion.

### Go Analysis

Gene Ontology (GO) analysis was performed using the topGO package in R. For both transcriptomic and proteomic datasets, the ontology was restricted to Biological Processes, and differentially significant genes were tested against all genes detected in the respective dataset.

Two statistical approaches were applied: a classic Fisher’s exact test and topGO’s weight01 algorithm. GO categories were considered statistically significant if the weight01 Fisher’s test yielded a p-value ≤ 0.05.

### Structure display

Protein structures were obtained from the Uniprot database as AlphaFold-predicted models and visualized using PyMOL (https://www.pymol.org/).

## DATA AVAILABILITY

The mass spectrometry proteomics data have been deposited to the ProteomeXchange Consortium via the PRIDE partner repository with accession numbers (PXD068586, PXD068589, PXD068593). All other data is available from the corresponding author on reasonable request.

## Acknowledgements

We thank Dr. Jixian Zhai and Dr. Jinbu Jia for providing the Arabidopsis thaliana increased intron retention datasets for various biotic and abiotic conditions. We also thank Timothy Pham for critically reading the manuscript. This work was funded by the National Institutes of Health grants R01GM135706 and S10OD030441 to S.-L.X. and diversity supplement to support A.V.R, and by the Carnegie Endowment Fund to the Carnegie Mass Spectrometry Facility.

## Author contributions

S.-L.X. conceptualization; A.V.R. and T.S.G. TMT sample preparation.

A.V.R. and C.Z. data analysis and figures. S.S.K., T.S.G., R.S., D.B., technical support. A.V.R. and S.-L.X. manuscript writing.

---

**Supplementary Figure S1.**
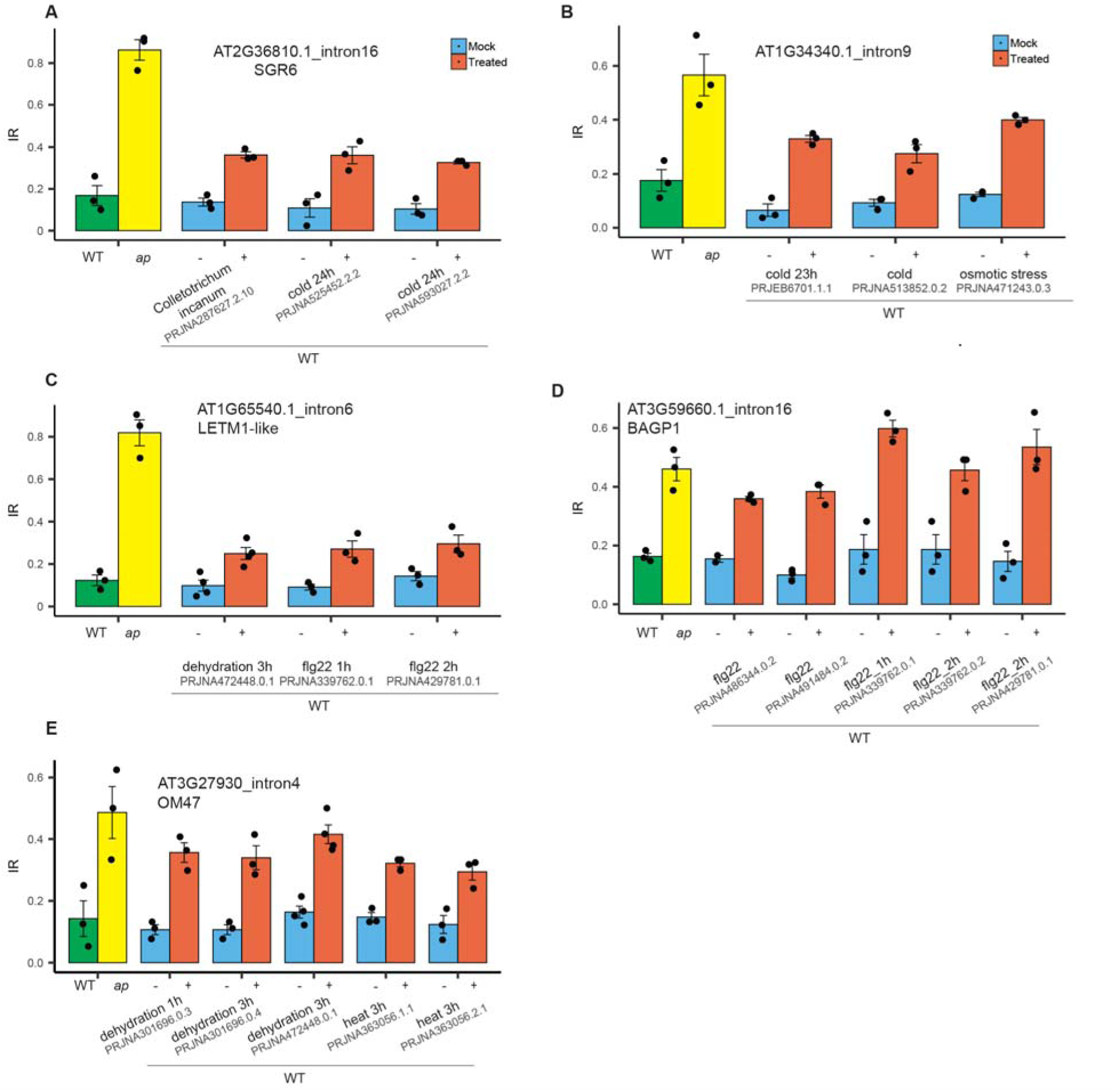
Intron retention (IR) events observed in the *acinus pinin* mutants are physiologically recapitulated in wild-type plants under biotic and abiotic stress conditions, as analyzed using *PlantIntronDB*. All IR events shown are significantly different from their respective controls (student t=test, p ≤ 0.05). (**A**-**E**) Bar graphs show five representative IR events altered in the *acinus pinin* mutant and also in wild-type plants following cold, osmotic, dehydration, flagellin, fungi, or heat treatment. The specific RNA-seq project names are indicated. Error bars represent the mean ± the standard deviation. The IR data for the *acinus pinin* (*ap*) mutant are from (44).

**Supplementary Figure S2.**
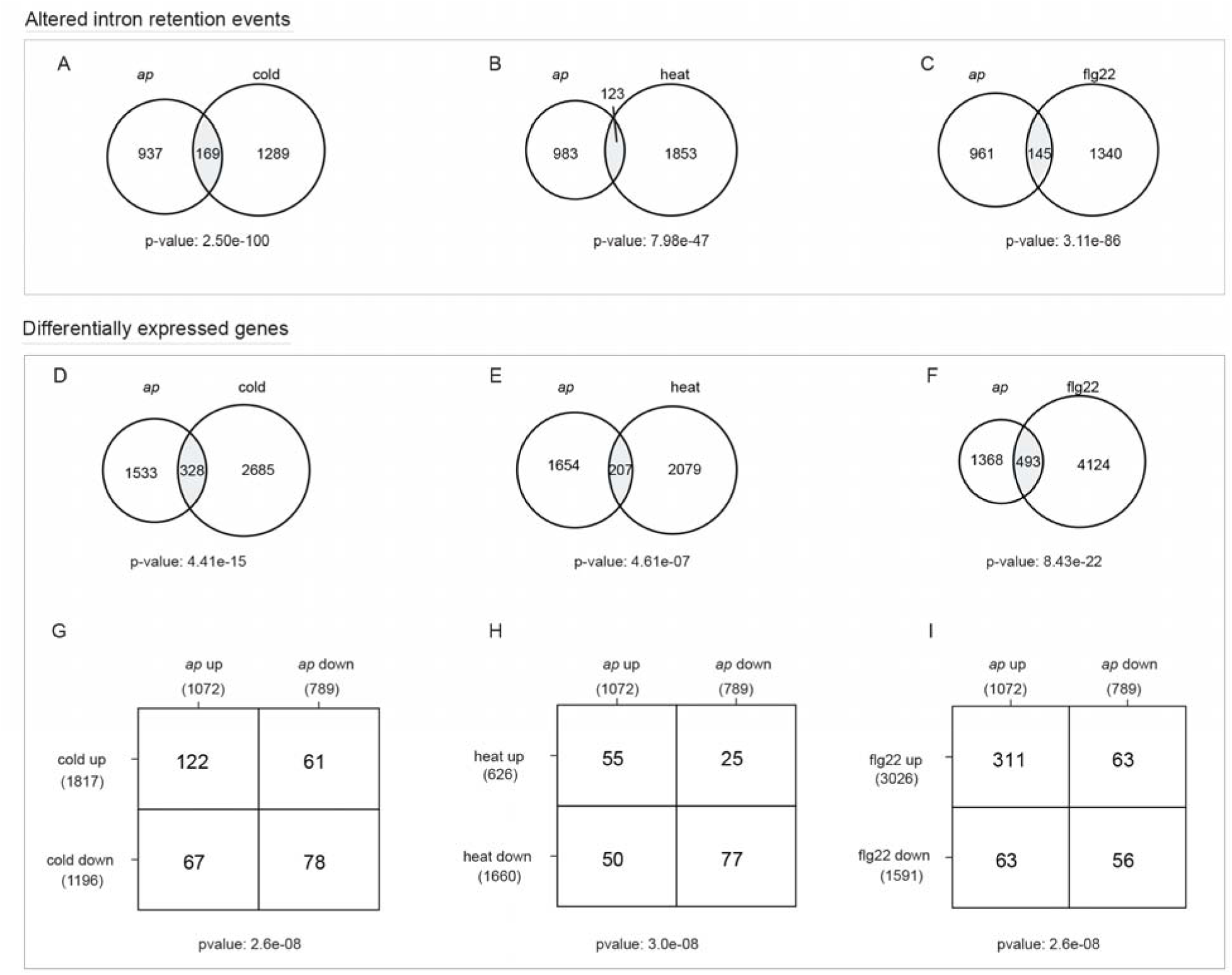
Overlap of altered intron retention (IR) events and differentially expressed genes (DEGs) in wild-type upon stress treatment and in the *acinus pinin* mutants. (**A**–**C**) Altered IR events overlap in wild-type with cold (A), heat (B), and flg22 (C) treatments and in *ap* mutants. (**D**–**F**) DEGs overlap in wild-type with cold (D), heat (E), and flg22 (F) treatments and in *ap* mutants. (**G**–**I**) Directional overlap of up-and down-regulated differentially expressed transcripts with cold (G), heat (H), and flg22 (I), classified as concordant (same direction) or discordant (opposite direction). Statistical significance was assessed using the hypergeometric test (A–F) and Fisher’s exact test (G–I).

**Supplementary Figure S3.**
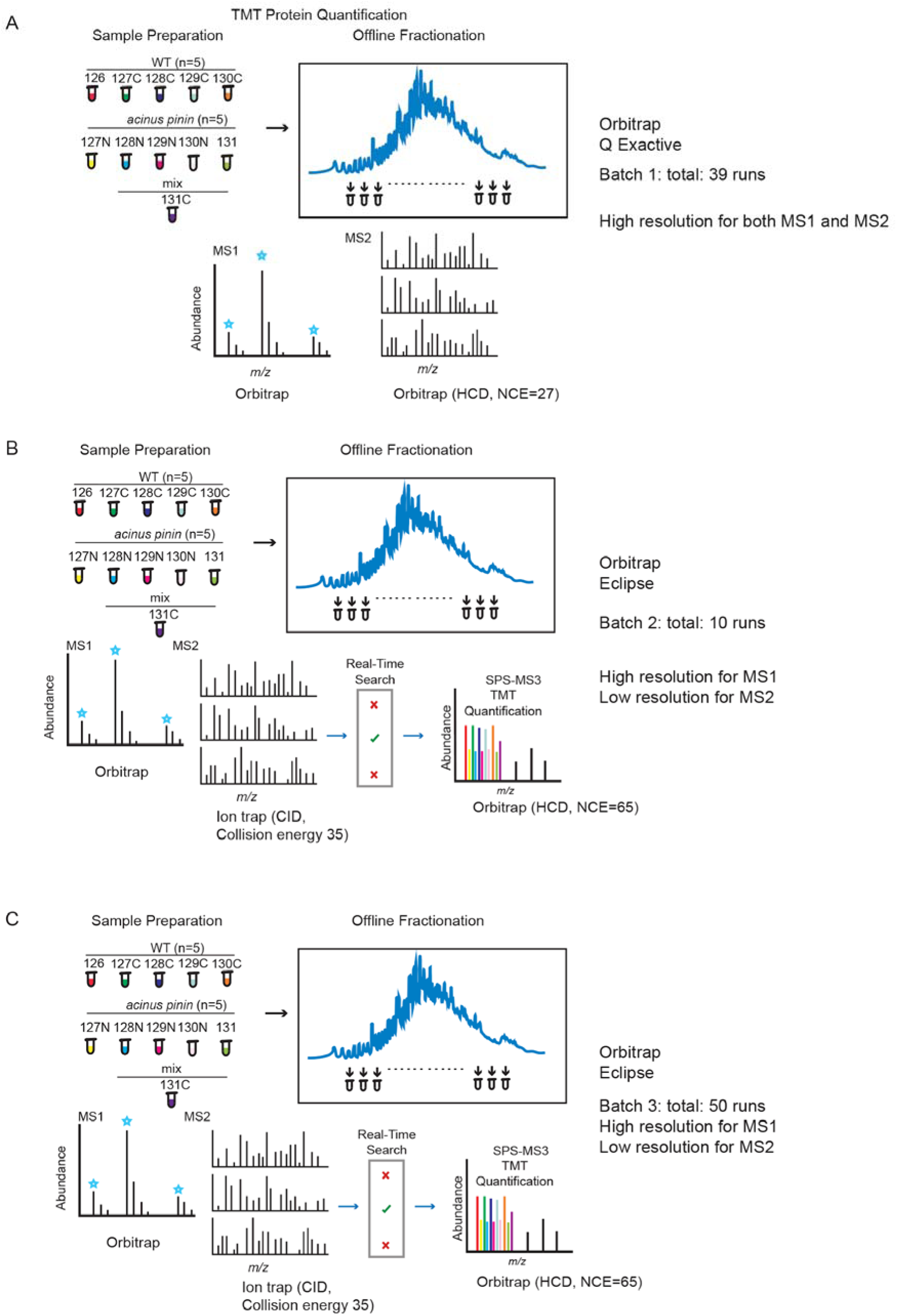
Tandem Mass Tag (TMT) quantitative analysis of wild-type and *acinus pinin* mutants. (**A**-**C**) Three batches of TMT experiments were performed: batch 1 on a Q Exactive with MS2-level quantification (panel A), and batches 2 and 3 on an Orbitrap Eclipse with MS3-level quantification (panels B and C). The two Orbitrap Eclipse experiments differ in fractionation numbers. Proteins from WT (5 replicates) and *acinus pinin* mutant (5 replicates) were trypsin-digested and labeled using TMT11-plex. The last channel was assigned to a pooled reference. The labeled peptides were combined and fractionated using high pH reverse-phase HPLC before being analyzed by LC-MS/MS. For the Orbitrap Eclipse, a narrow 0.7 Da precursor isolation window and a real-time library search (RTS)-SPS-MS3 were applied to enhance sensitivity and precision by targeting library-matched TMT peptides and synchronously isolating MS2 fragments for MS3 fragmentation (73).

**Supplementary Figure S4.**
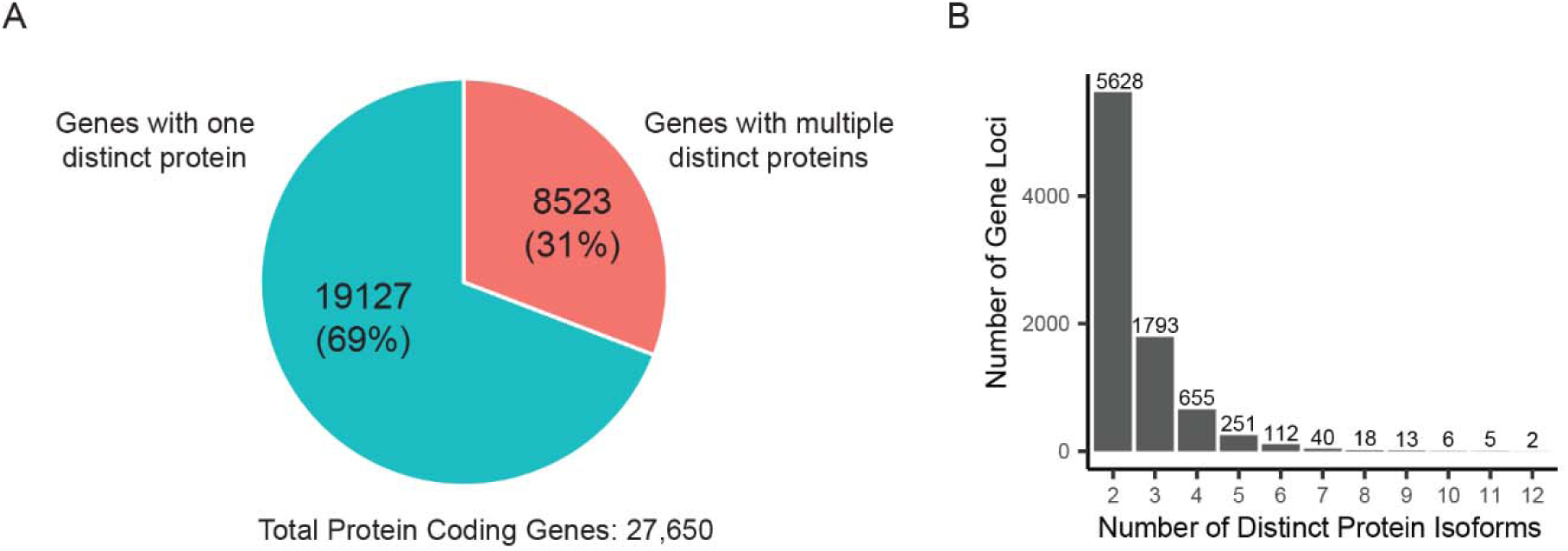
Prevalence and distribution of alternative isoforms in the Araport11 annotation. **(A)** Proportion of genes predicted to produce a single isoform versus multiple isoforms. Approximately 31% of genes are annotated with more than one isoform protein. Genes that have different transcript isoforms but produce only one unique protein or similar protein sequence are collapsed. **(B)** Distribution of the number of protein isoforms among genes with multiple isoforms in Araport11 database.

**Supplementary Figure S5.**
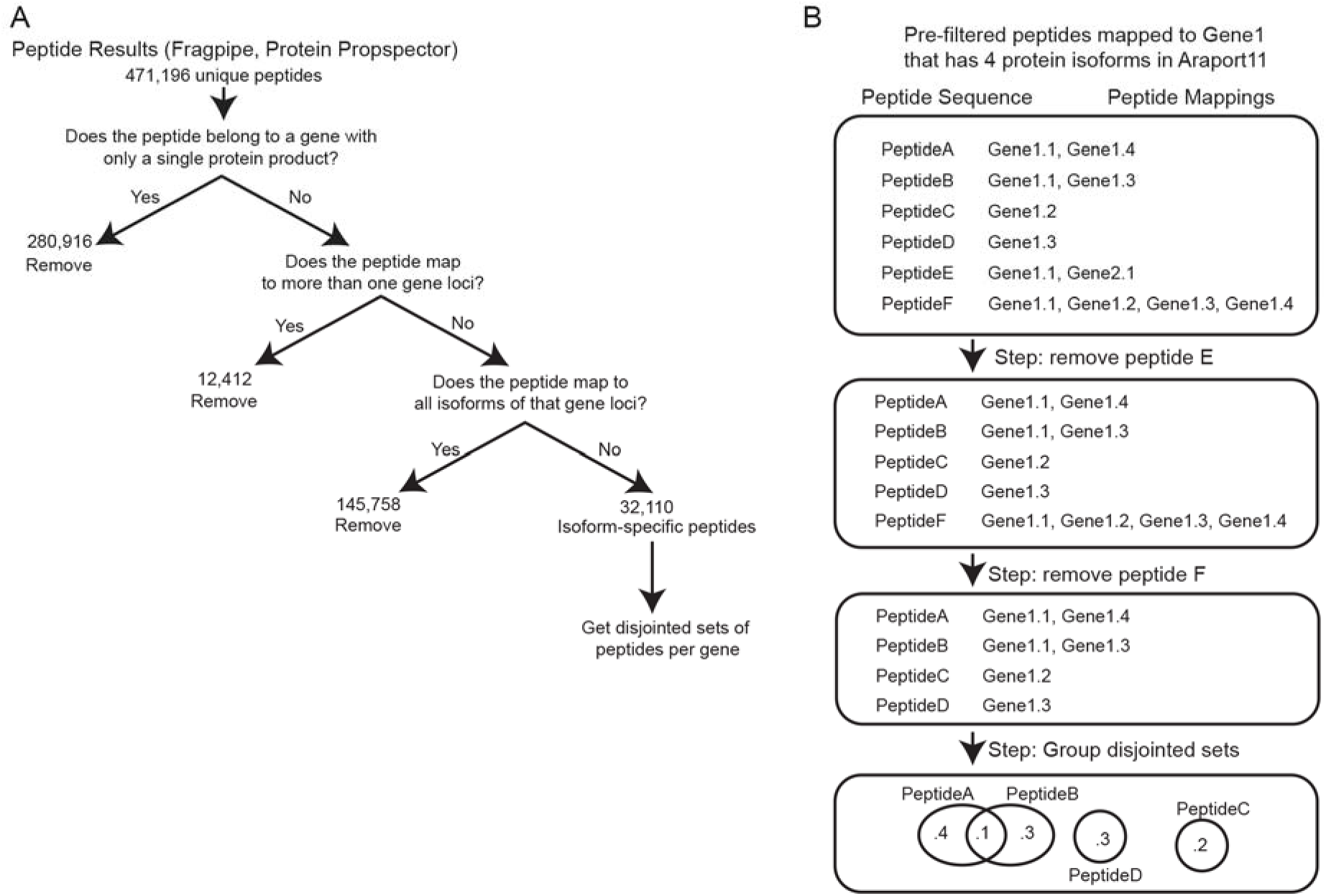
Workflow for protein isoform calculation. (**A**) Summary of the filtering strategy. A total of 471,196 peptides were initially identified from total datasets; after filtering, 32,110 isoform-specific peptides remained for isoform mapping. (**B**) Representative cartoon illustrating how specific peptides are excluded from isoform calculations based on defined criteria.

**Supplementary Figure S6.**
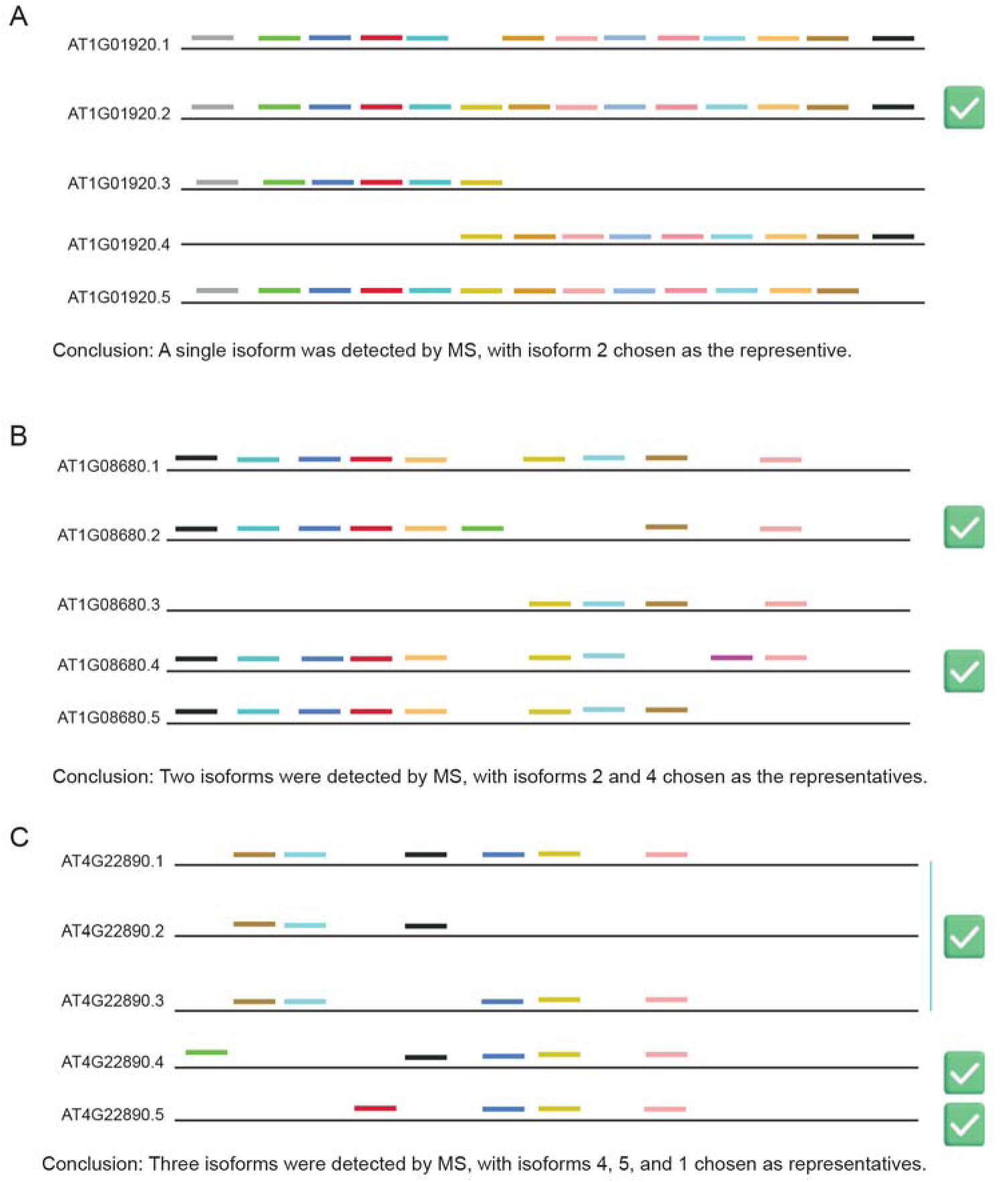
Peptide-based isoform analysis for genes with multiple annotated isoforms. (**A**–**C**) Schematics illustrating peptide assignments and isoform calculations for AT1G01920, AT1G08680, and AT4G22890. Calculations are based on a hypothesized predominant isoform, defined as the one supported by the greatest number of isoform-specific peptides. This provides a conservative estimate; the actual number of isoforms may be higher. Colored lines represents unique peptides aligned to each predicted isoform.

**Supplementary Figure S7.**
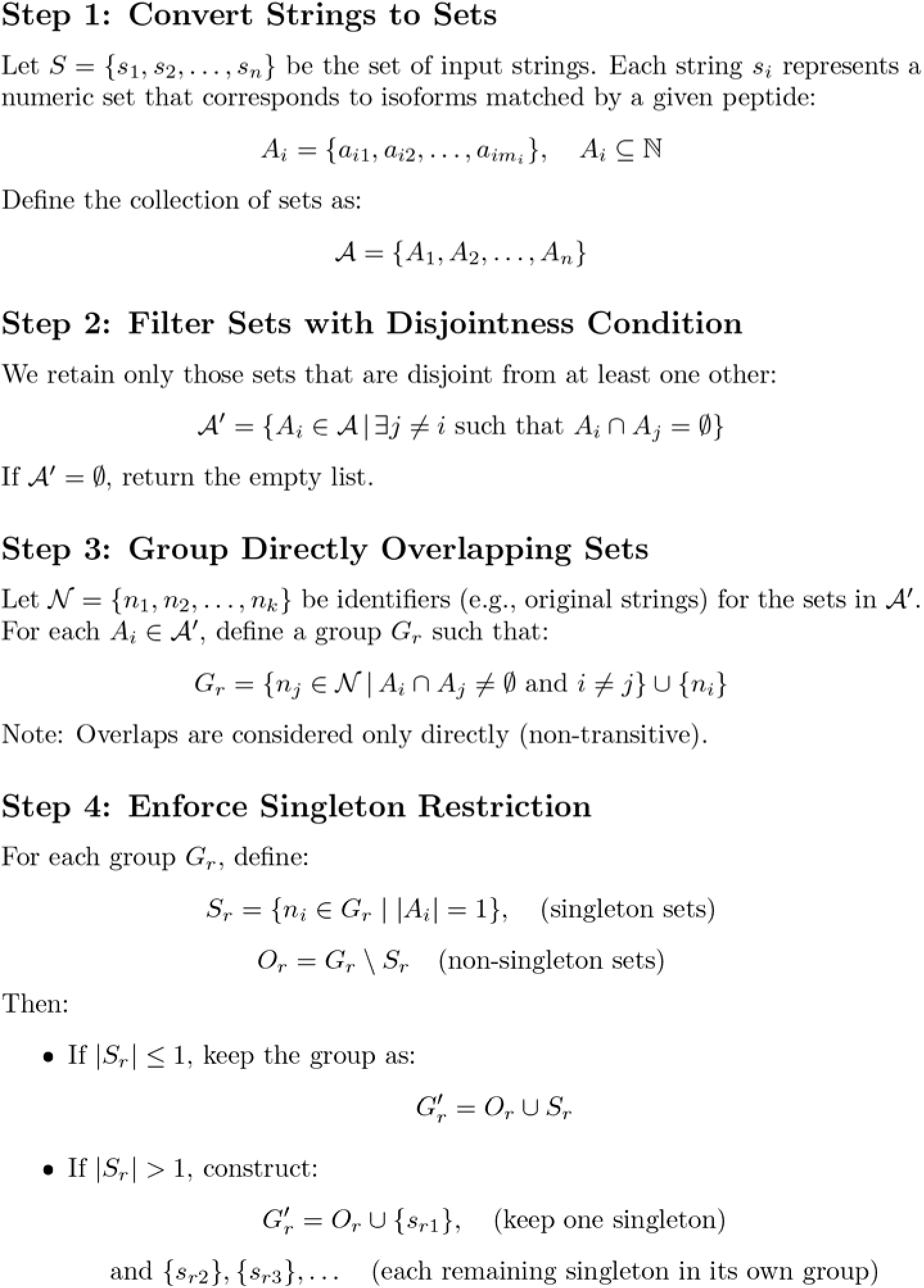
Workflow for filtering and grouping numeric sets. Numeric sets represent isoforms mapped by each peptide. (1) Convert peptide mappings into numeric sets. (2) Retain sets disjoint from at least one other set. (3) Group sets with direct overlaps. (4) Retain only one singleton per group; additional singletons are reassigned to separate groups. This figure was generated by AI based on the R code used in our isoform calculation algorithm.

**Supplementary Figure S8.**
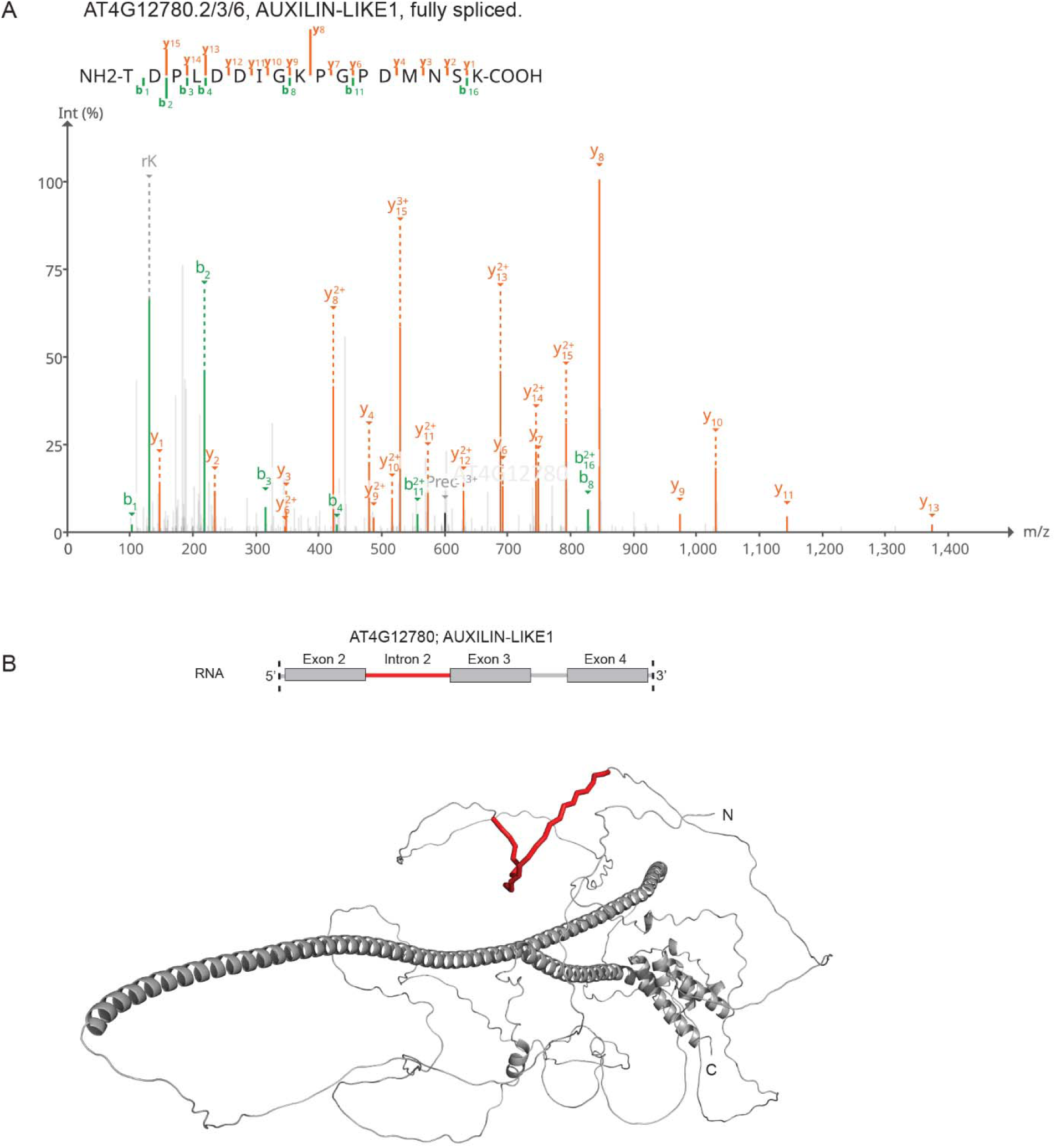
Spectrum of a junction peptide supporting the spliced form of AT4G12780, and the AlphaFold-predicted structure illustrating the location of the additional sequence resulting from intron retention. **(A)** MS2 spectrum of a peptide spanning the junction between exons 2 and 3 of *AT4G12780.2/3/6*, supporting the fully spliced Auxilin-like 1 isoform. The spectrum exhibits extensive b-and y-ion coverage, unambiguously identifying exon 2–3 junction peptide sequence. **(B)** AlphaFold-predicted structure of AT4G12780, in which the retained intronic sequence is highlighted in red. The additional 27 amino acids derived from intron retention are incorporated into a predicted unstructured region based on AlphaFold model ID: AF-Q9SU08-F1.

**Supplementary Figure S9.**
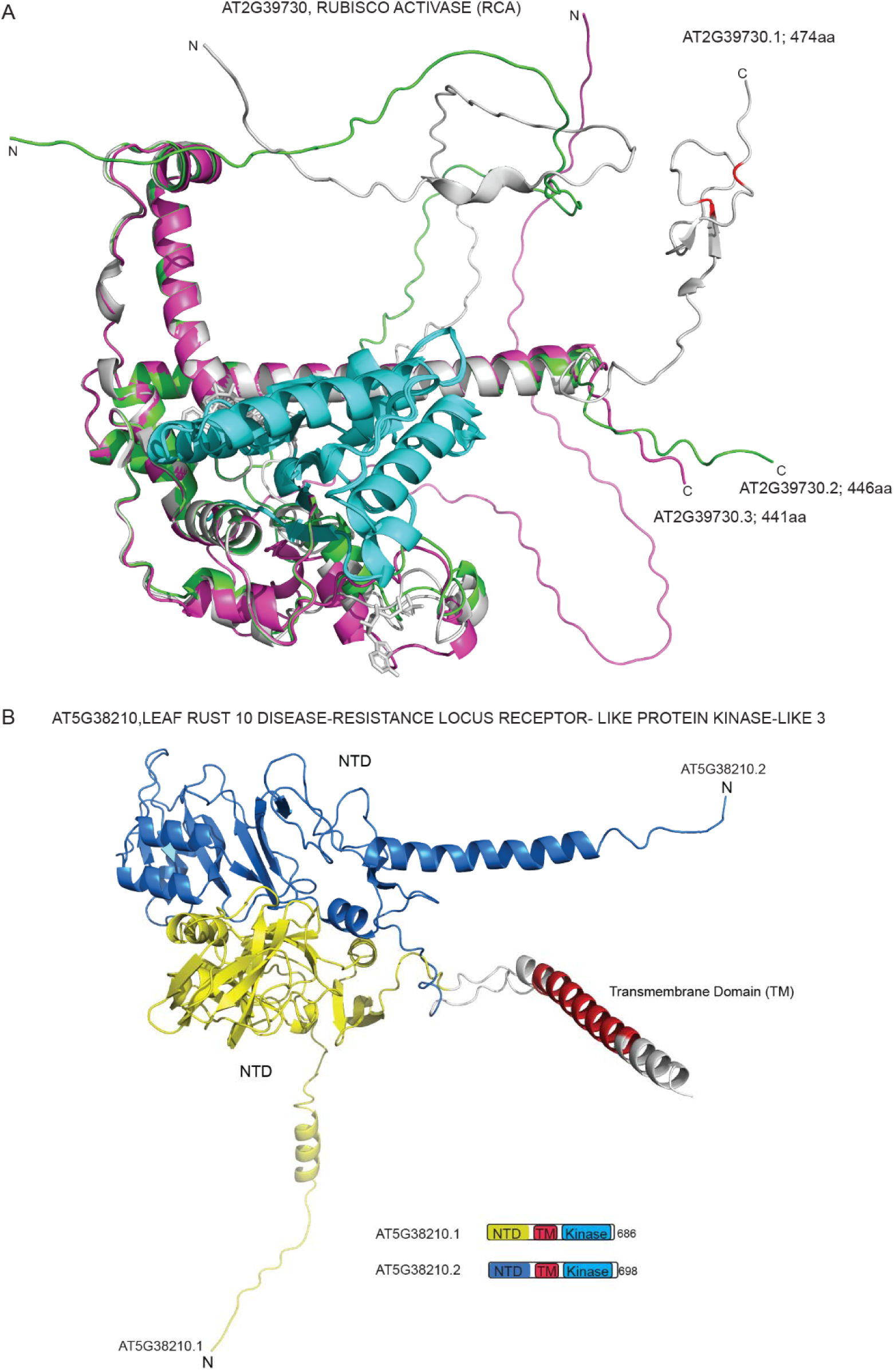
Structure information for protein isoforms of RCA and LRK10L3. **(A)** AlphaFold-predicted structures of three RCA isoforms (UniProt ID: P10896) showing a shared N-terminal region but variable C-terminal domains. The full-length isoforms contain two C-terminal cysteine residues highlighted in red, which are implicated in redox regulation. **(B)** AlphaFold-predicted structures of two LRK10L3 isoforms (AlphaFold model IDs: AF-A0A1P8BDK9-F1 and AF-Q8VYG0-F1) showing distinct N-terminal regions and a shared transmembrane (TM) domain. The C-terminal kinase domain is omitted for clarity.

**Supplementary Figure S10.**
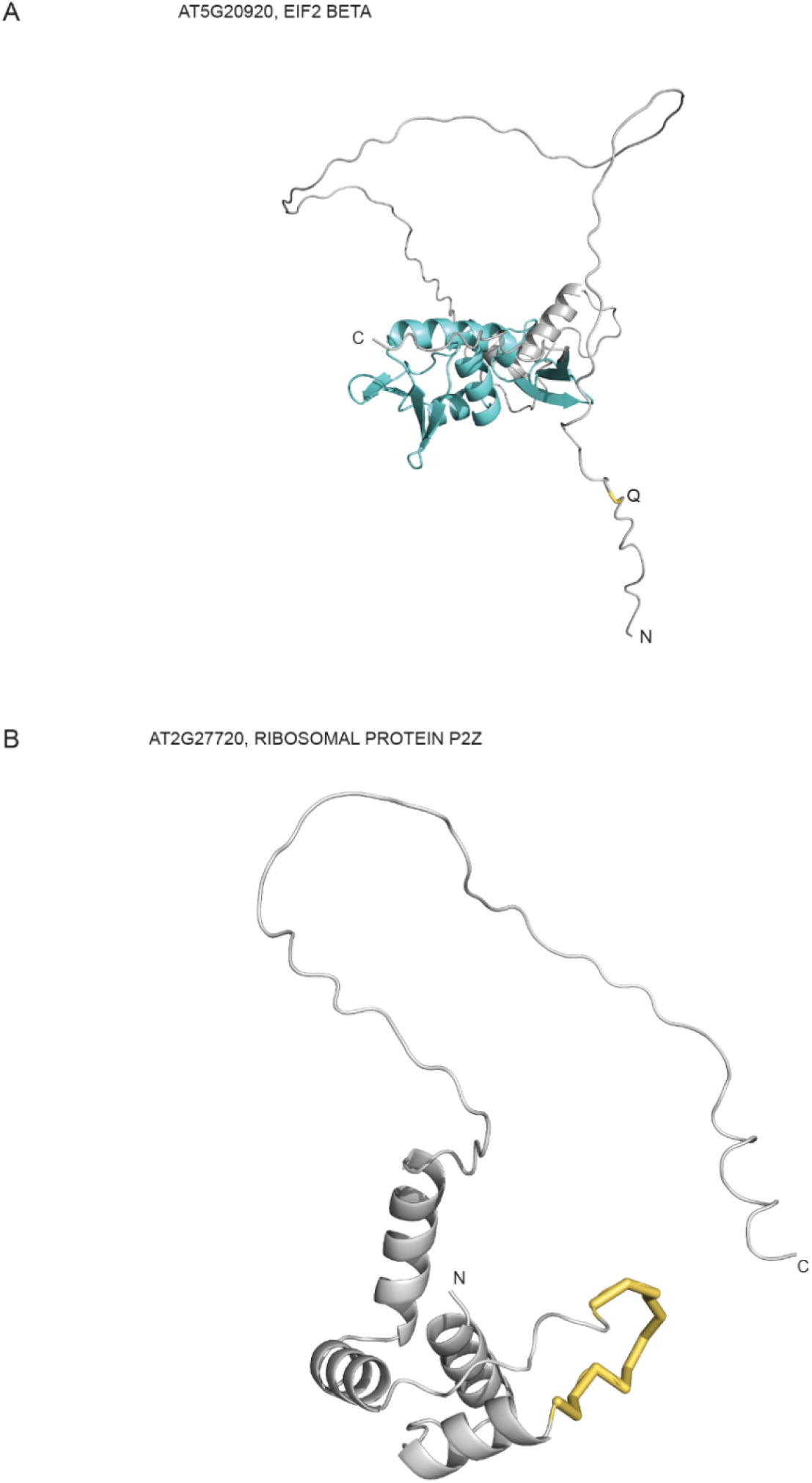
Structure information for EIF2 BETA and RIBOSOMAL PROTEIN P2Z. (**A**) AlphaFold-predicted structure of EIF2 BETA (AF-Q41969-F1-model_v6), with the glutamine (Q) residue added by alternative splicing highlighted. (**B**) AlphaFold-predicted structures of ribosomal protein P2Z (AlphaFold model IDs: AF-A0A1P8BDK9-F1 and AF-Q8VYG0-F1), showing the additional 12-amino-acid sequence difference (no exon3 skipping) highlighted in yellow.

**Supplementary Figure S11.**
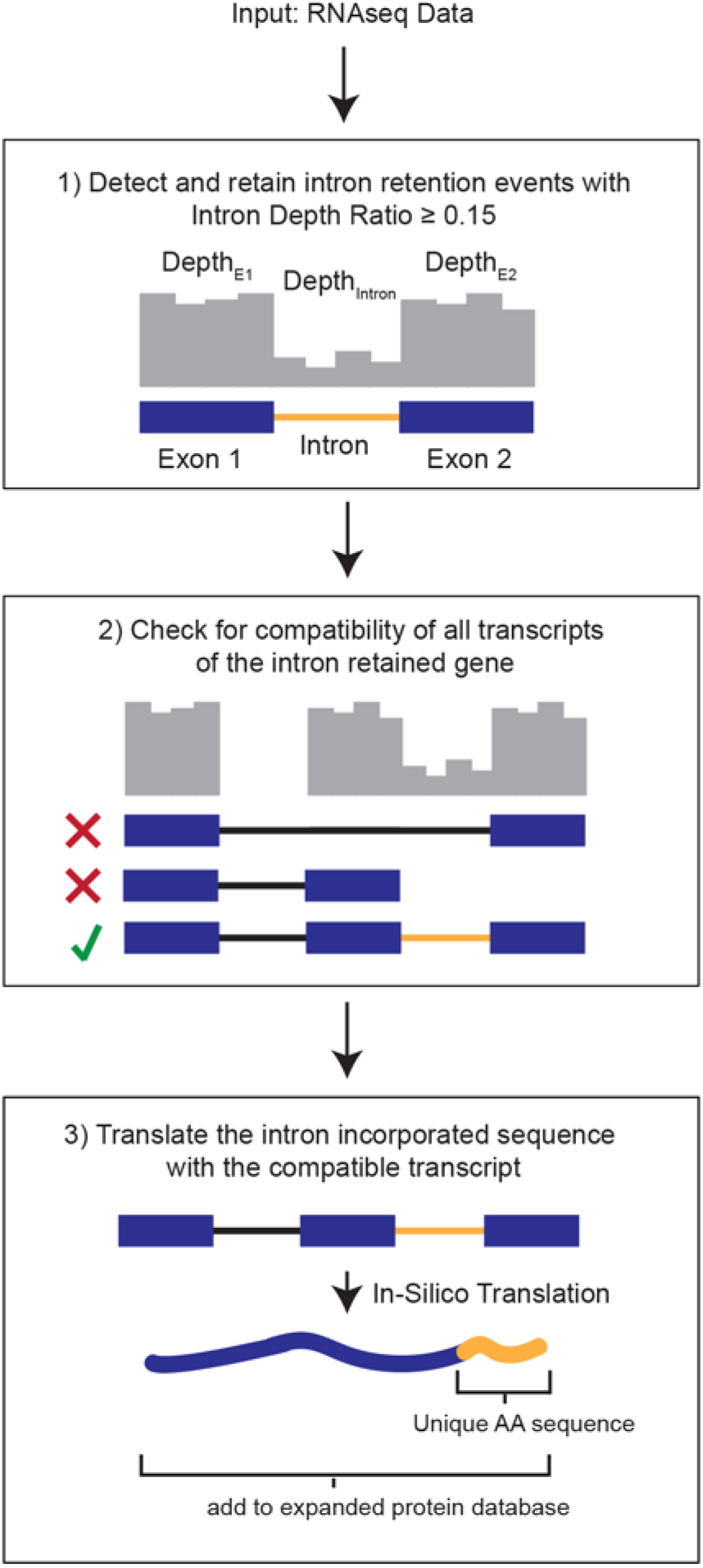
Workflow for generating a custom peptide database from transcripts containing unannotated retained introns. This multi-step pipeline generates novel peptide sequences from transcripts with retained introns identified via RNA-seq. (1) Intron retention events are filtered to exclude low-confidence cases (Intron Depth Ratio < 0.15). (2) Retentions compatible with annotated exon boundaries in Araport11 are retained (i.e., flanked by two exons within the same transcript). Because RackJ collapses transcripts by gene loci before defining introns, intron boundaries defined by RackJ are not always directly compatible with the original transcript structure. Step 2 ensures that each intron is correctly integrated into its transcript. (3) Retained introns are incorporated into modified transcripts, translated *in silico* to produce protein sequences, and appended to the Araport11 proteome for downstream database searches.

**Supplementary Figure S12.**
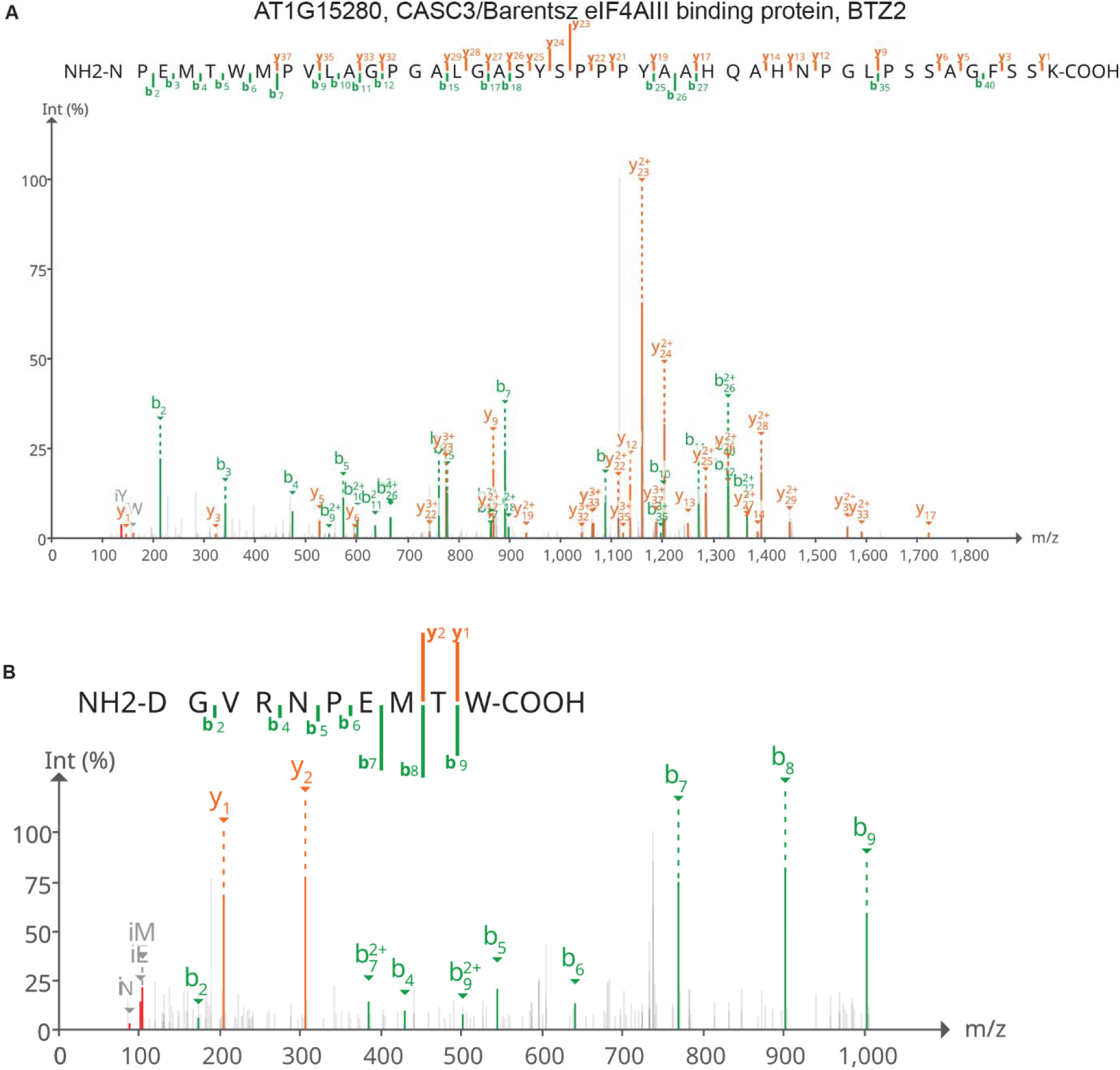
MS2 spectra of peptides supporting both the spliced and intron-retained isoforms of AT1G15280 (BTZ2); the intron-retained isoform is not annotated in Araport11. **(A)** Peptide spanning the exon 6–exon 7 junction of the spliced isoform AT1G15280.1. **(B)** Peptide spanning the exon 6–intron 6 junction of the intron-retained isoform. The peptide was detected in both WT and *acinus pinin* mutants.

**Supplementary Figure S13.**
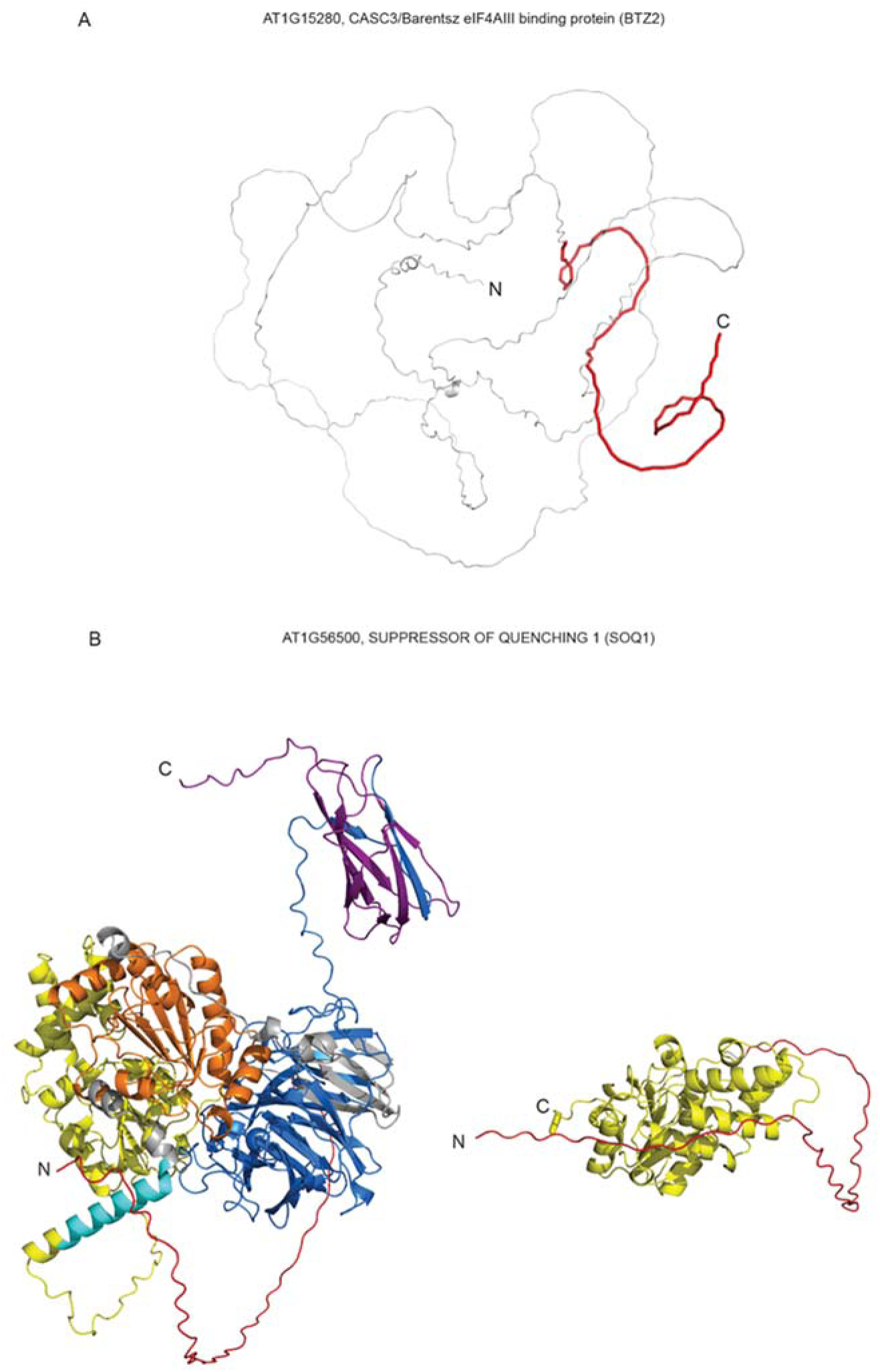
Structural information for BTZ2 and SOQ1. (**A**) AlphaFold-predicted structure of BTZ2, showing that the protein is largely disordered. Intron retention of intron 6 introduces a premature stop codon, resulting in a truncated protein that is 84 amino acids shorter. The missing region is highlighted in red. (**B**) Structural comparison of the full-length (left) and truncated (right) forms of SOQ1. The color scheme is consistent with that in Figure 4F.

**Supplementary Figure S14.**
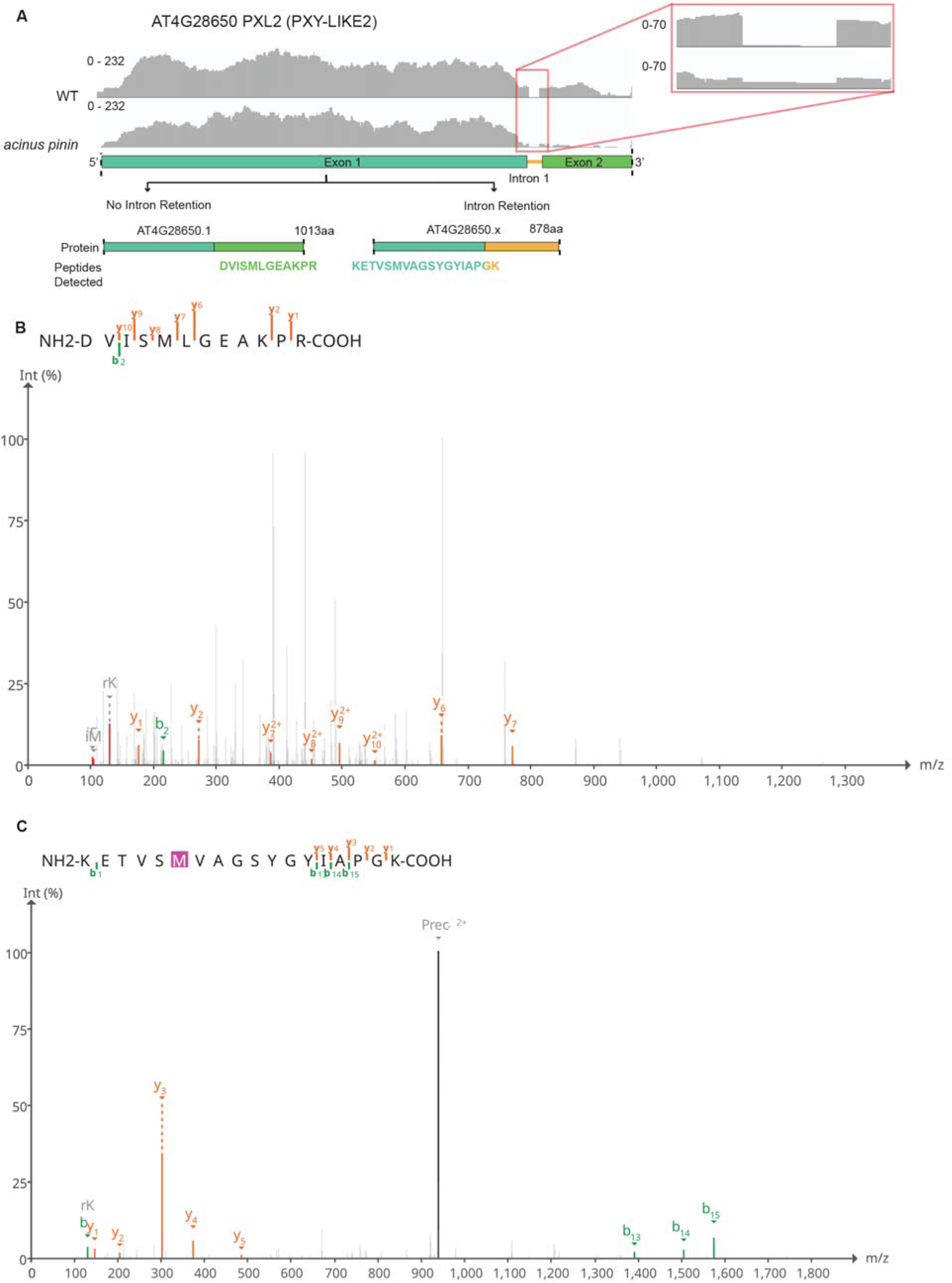
Proteomic evidence for two isoforms of AT4G28650 (PXY-LIKE 2): the annotated spliced form and an intron-retained isoform absent from Araport11. (A) RNA-seq coverage and gene annotation showing retention of intron 1 in both WT and *acinus pinin*. Inset highlights reads spanning intron 1, revealing increased retention in the *acinus pinin* mutant. (B) MS2 spectrum detecting the peptide encoded by exon 2, supporting the canonical spliced isoform (AT4G28650.1). (C) MS2 spectrum detecting the junction peptide spanning exon 1 and intron 1, supporting a previously unannotated intron-retained isoform (AT4G28650.x). Retention of intron 1 introduces a stop codon, resulting in a truncated protein of 878 amino acids. This junction peptide was detected in multiple tissues across cell atlas datasets.

**Supplementary Figure S15.**
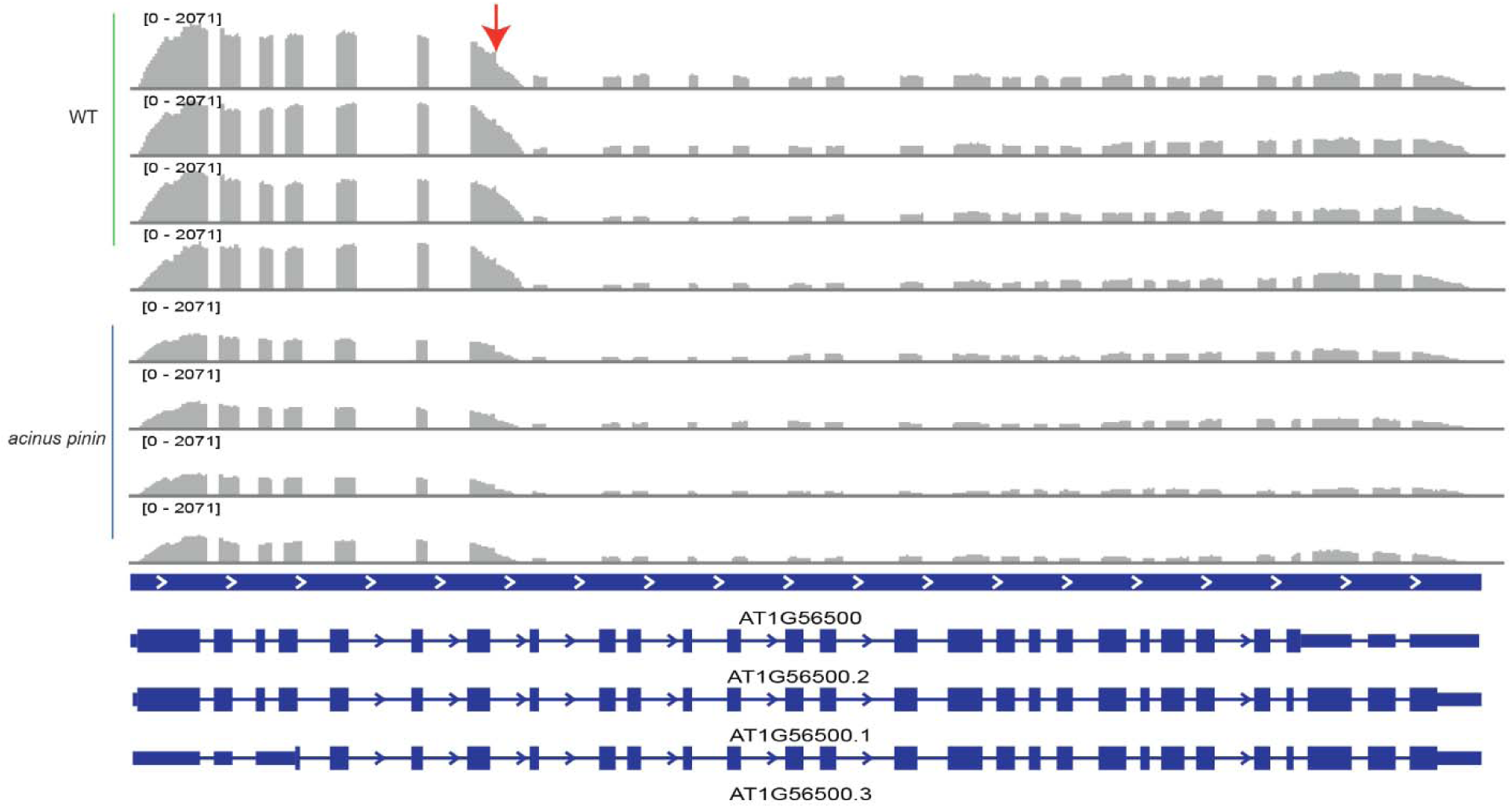
RNA-seq coverage indicates the need for a revised gene model of AT1G56500. RNA-seq results and schematic annotation of canonical AT1G56500 isoforms are shown. Coverage from four biological replicates of WT and *acinus pinin* shows consistent retention of intron 7, accompanied by a sharp drop in downstream reads. This pattern suggests that, in addition to the current annotation (AT1G56500.1/2/3), a shorter mRNA isoform is generated via intron 7 retention and alternative 3’ processing within intron 7, with transcript levels reduced in the *acinus pinin* mutant.

**Supplementary Figure S16.**
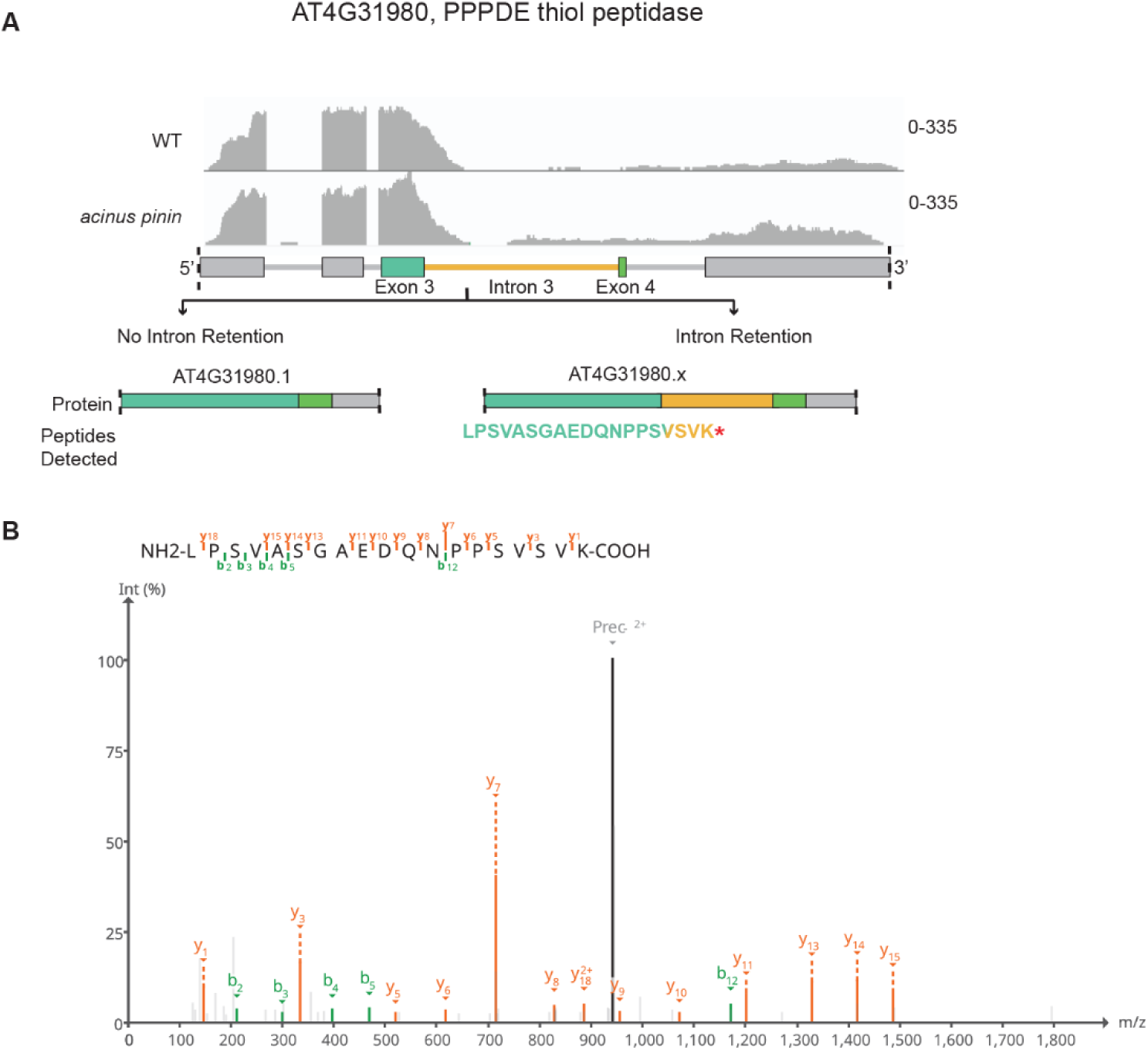
Proteomic and transcriptomic analyses reveal a new isoform of AT4G31980 (PPPDE thiol peptidase), which includes intron 3. (A) RNA-seq coverage is compared with the Araport11 annotation. The reads demonstrate robust coverage across exons 1–3 and the 5′ region of intron 3. There is a gradual decline in read coverage across the remainer of the intron and downstream regions. This suggests that the current gene model requires refinement. However, expression of the annotated isoform AT4G31980.1 cannot be ruled out based on the 3’ reads of the gene. Several AT4G31980 peptides were identified; however, peptides supporting the junction of exons 3 and 4 were not detected. (B) The MS2 spectrum identifies a peptide that spans exon 3 and intron 3. The extensive b-and y-ion series support unambiguous identification of the peptide.

**Supplementary Figure S17.**
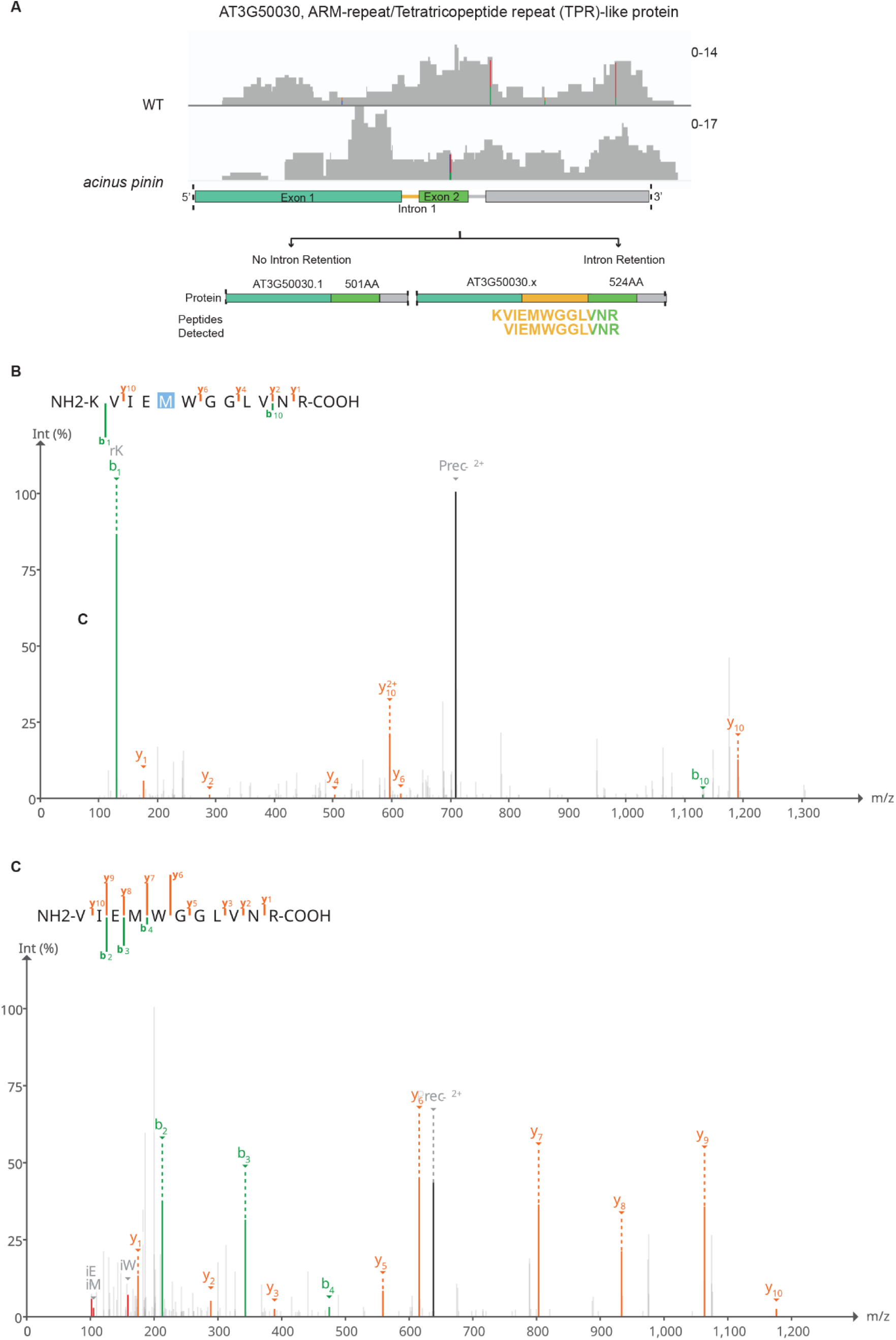
Proteomic evidence supports translation of annotated intron 2 in AT3G50030 (ARM-like/tetratricopeptide repeat protein), suggesting a revised isoform. (**A**) RNA-seq coverage and gene annotations from Araport11 (AT3G50030.1) and the proposed revised model (AT3G50030.x). The revised model includes intron 1, although low RNA-seq coverage makes precise boundary determination challenging. Intron 1 is in-frame and lacks a stop codon. (**B**–**C**) Two peptides were identified that map to intron 1, supporting its translation and indicating that the Araport11 annotation may require revision. Although TAIR currently lists AT3G50030 as “not expressed in wild-type plants,” our proteomic data detect multiple unique peptides from this protein, whereas no junction peptides were found to support AT3G50030.1.

**Supplementary Figure S18.**
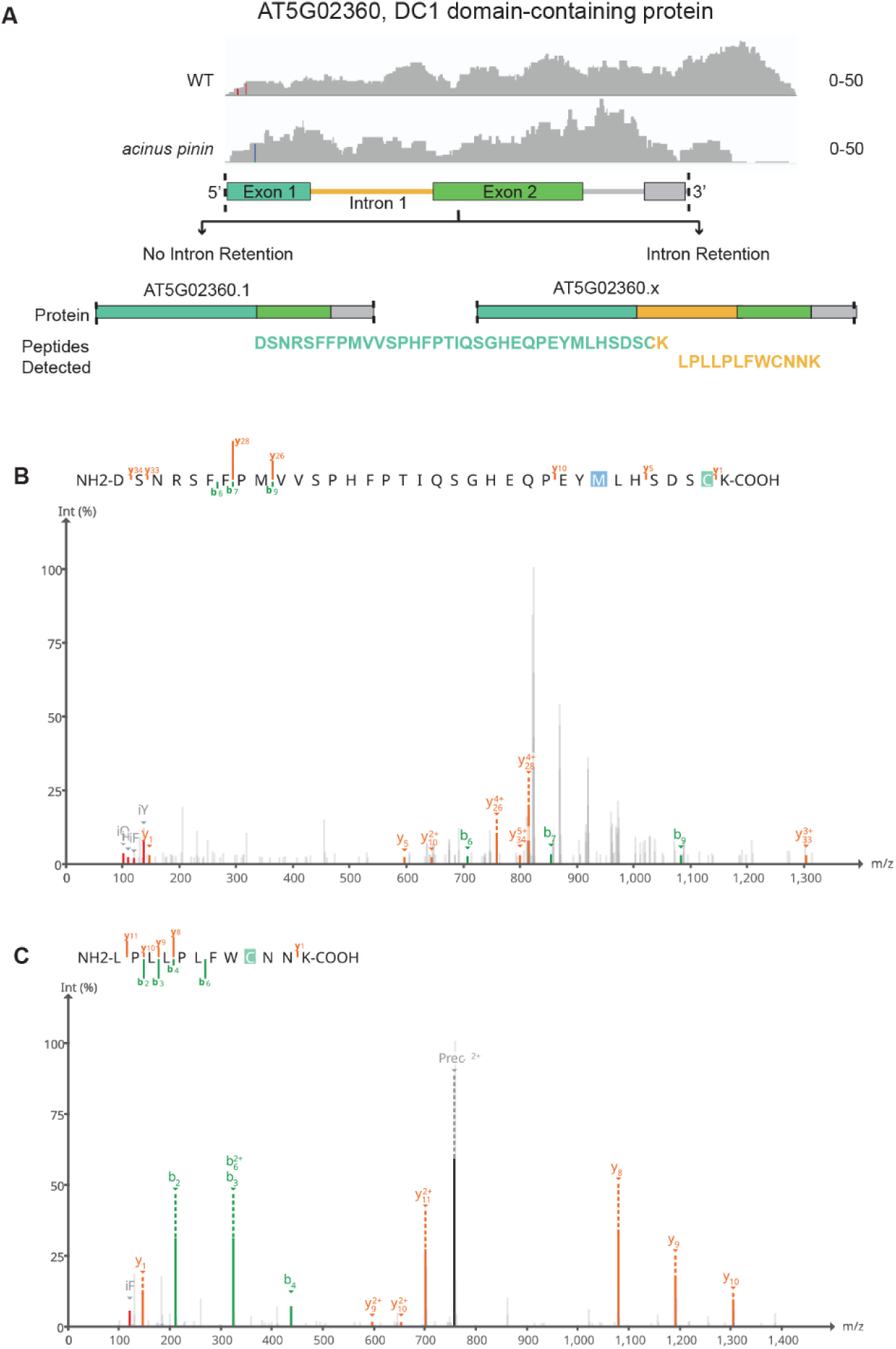
Proteomic evidence supports a revised annotation of AT5G02360 (DC1 domain-containing protein) compared with the current annotation (AT5G02360.1). (**A**) RNA-seq coverage and Araport11 annotation for AT5G02360. Although RackJ interprets these reads as intron retention based on the existing annotation, the low RNA-seq coverage makes it difficult to precisely define intron boundaries. (**B**) Two independent MS2 spectra provide peptide-level evidence supporting a revised annotation in which intron 1 is retained and translated. Multiple peptides were detected for AT5G02360, but no isoform-specific peptides were identified for AT5G02360.1.

**Supplementary Figure S19.**
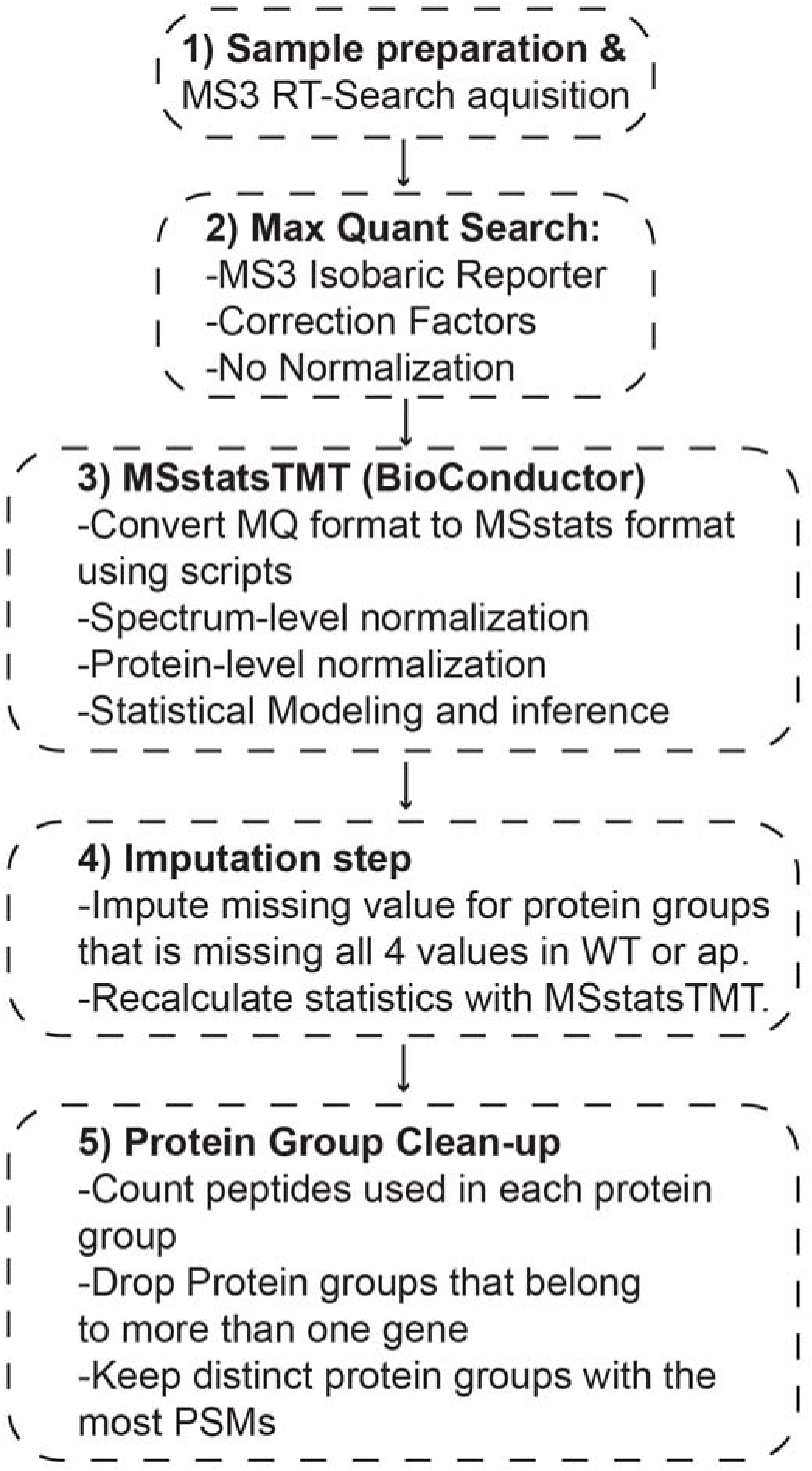
: Workflow for protein-level quantification using MS3 real-time library search data. Raw MS3 data were processed in MaxQuant with the search type set to MS3 Isobaric Reporter. Correction factors were applied as appropriate, and normalization within MaxQuant was omitted. MSstatsTMT was then used for normalization and statistical analysis. To generate a single quantification value per protein, protein groups were filtered to retain only those uniquely mapping to a single gene. When multiple protein groups corresponded to the same gene, the group with the highest number of detected peptides was selected.

**Supplementary Figure S20.**
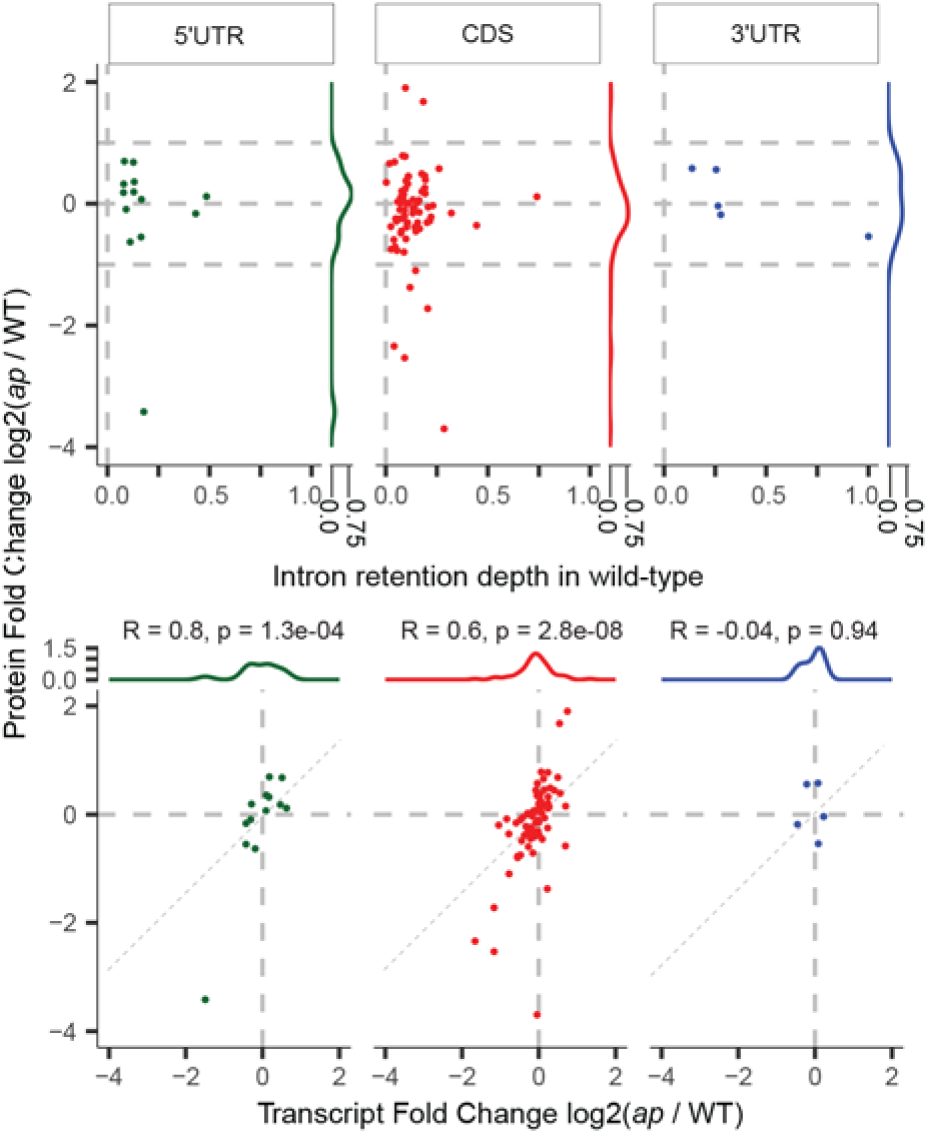
Effects of retained introns that are reduced in *acinus pinin* mutants on transcript and protein abundance. Density plots show distribution changes in protein (top) and RNA (bottom) levels for genes with retained introns that are reduced in *acinus pinin* mutants. Intron retention depth in wild-type is shown in the top panel. Interpretation is limited due to the small number of events and potential confounding effects from increased retention of other introns in the same genes in the mutant.

## Notes

### Competing Interest Statement

The authors have declared no competing interest.

### Summary of Updates

Revised the writing and several supplemental figures.

